# Dissociable effects of subliminal incentives on initiation and velocity control of goal-directed eye movements

**DOI:** 10.1101/2020.11.11.377986

**Authors:** Vasko Kilian Hinze, Ozge Uslu, Jessica Emily Antono, Melanie Wilke, Arezoo Pooresmaeili

**Author notes:** VKH and OU are joint first authors. MW and AP are joint last authors.

## Abstract

Over the last decades, several studies have demonstrated that conscious and unconscious reward incentives both affect performance in physical and cognitive tasks, suggesting that goal-pursuit can arise from an unconscious will. Whether the planning of goal-directed saccadic eye movements during an effortful task can also be affected by subliminal reward cues has not been systematically investigated. We employed a novel task where participants had to make several eye movements back and forth between a fixation point and a number of peripheral targets. The total number of targets visited by the eyes in a fixed amount of time determined participants’ monetary gain. The magnitude of the reward at stake was briefly shown at the beginning of each trial and was masked by pattern images superimposed in time. We found that when reward cues were fully visible and thus consciously perceived, higher reward enhanced all saccade parameters. However, a dissociation was observed between the effects of subliminal rewards on saccade initiation and peak velocities. While truly subliminal reward cues did increase the number of saccades, they did not enhance saccades’ peak velocity. Additionally, participants who had reached a truly subliminal level of reward perception showed a decrement in accuracy as a function of reward across all visibility levels, as saccade endpoint error was larger when higher reward incentives were expected. This suboptimal speed-accuracy trade-off did not occur in the supraliminal group. These results suggest that although saccades’ initiation can be triggered by subconscious mechanisms, conscious awareness is required to optimally adjust the velocity and accuracy of eye movements based on the expected rewards.

## Introduction

Human behavior is guided by a drive to maximize survival chances through avoiding unwanted situations and approaching desirable outcomes, i.e. rewards (Pessoa and Engelmann, 2010). However, human’s cognitive or physical resources, for instance the extent to which they can exert physical effort in a task in order to obtain a reward, are limited. Hence, in order to efficiently interact with the surroundings, in a way that rewards are maximized while the effort to attain them is kept as low as possible, agents often set a certain goals at a given point in time and adjust their performance accordingly (Dickinson and Balleine, 1994). A longstanding idea in cognitive neuroscience posited that the underlying processes of goal-pursuit arise from conscious awareness, where an agent is aware of the content of what one is experiencing or trying to achieve and the costs entailed (Kouider and Dehaene, 2007; Zedelius et al., 2014). In the last decades, this idea has been challenged as researchers have repeatedly shown that goal-pursuit can also have its origin in an unconscious mind or even operate without conscious awareness (Bargh et al., 2001; Bijleveld et al., 2012; Custers and Aarts, 2005; Hart and Albarracín, 2009; Pessiglione et al., 2007). Here, we ask whether these effects also extend to goal-directed planning of eye movements.

Our study is inspired by the pioneering work of Pessiglione et al. (Pessiglione et al., 2007) investigating whether goal-pursuit could be influenced by subliminally presented reward incentives. It was demonstrated that participants can adjust their level of physical effort in a handgrip force task dependent on the magnitude of the reward at stake in each trial. Remarkably, monetary reward incentives not only increased the exerted force when they were consciously perceived (i.e. were supraliminal) but also invigorated performance when presented below the conscious threshold (i.e. subliminally). Likewise, the analysis of the simultaneously acquired fMRI data demonstrated that the same subcortical brain structures encode consciously as well as subconsciously perceived reward incentives.

A number of subsequent studies replicated these results extending them to other behavioural paradigms and mental versus physical effort. A study using a memorization task showed an increase of performance for both supra- and subliminally presented high reward cues (Bijleveld et al., 2009). Similarly, the invested effort in a mental arithmetic task (Bijleveld et al., 2010) or a task that involved updating memorized digits by applying an arithmetic operation to them (Capa et al., 2011) was found to be enhanced by both consciously as well as subconsciously perceived rewards. The increased effort could show itself in the form of increased speed or accuracy (Bijleveld et al., 2012, 2009), which are commonly used measures in evaluating performance (Zedelius et al., 2014). Together, these findings indicate that goal-pursuit is not only affected by consciously perceived incentives but could also be steered by unconscious mechanisms.

Nevertheless, differences between the effects of conscious and unconscious reward incentives have also been reported. In a recent study it was demonstrated that although both supraliminal and subliminal reward cues produced the same behavioural outcome, the underlying neural dynamics seemed to be vastly different (Bijleveld et al., 2014). Furthermore, other studies showed that unconscious reward processing is rather limited when it comes to improving performance strategy and efficiency during complex task contexts (Bijleveld et al., 2010; Zedelius et al., 2012). Thus, reward seems to show divergent effects when perceived consciously or unconsciously in more complex tasks.

In the current study, we ask whether humans will undertake a cost-benefit adjustment of their oculomotor effort based on the reward incentives and whether this effect occurs even when reward cues are presented subliminally. Reward effects on eye movements are ubiquitous, as reported by a number of previous studies (for a review see Trommershäuser et al., 2009). For instance, neurophysiological studies in monkeys have demonstrated that the time taken to initiate a stimulus driven saccade (i.e. saccade latency) is significantly shorter when a higher reward is expected (Hikosaka, 2007; Hikosaka et al., 2000; Itoh, 2002). Furthermore, in rewarded conditions saccades typically have higher peak velocities [19, see also 20]. Importantly, the reward-related increase in saccades’ peak velocity cannot be fully explained by the stereotypic relationship between saccades’ velocity and amplitude known as *main sequence* (Bahill et al., 1975), demonstrating that rewards incentives affect saccades’ velocity above and beyond biomechanical factors. Similar results were found in human psychophysics studies as monetary rewards were shown to increase saccade velocity (Chen et al., 2014) and vigor (i.e. the peak velocity as a function of amplitude) (Choi et al., 2014; Reppert et al., 2015; Shadmehr et al., 2019). Interestingly, the increase in the speed of the saccade is not associated with a decrease in their accuracy (speed-accuracy trade-off), as rewards seem to increase accuracy and speed at the same time without having to give in to speed accuracy tradeoff (Manohar et al., 2015). Multiple lines of evidence suggest that modulation of saccade parameters by reward value is the result of a tight interaction between the neuronal underpinnings of reward processing and saccade planning (Hikosaka et al., 2014, for reviews see 2000; McCoy and Platt, 2005), at subcortical and cortical levels.

Despite the wealth of studies on the impact of rewards on different saccade parameters, it is not known whether humans can voluntarily control their saccadic *effort* based on the magnitude of the reward incentives (but see Muhammed et al., 2020 for evidence on voluntary albeit instructed control of the saccade velocity). Likewise, it is not known whether a putative reward-related adjustment of saccadic effort also occurs when the incentives are perceived subliminally, as has been the case for tasks requiring manual or cognitive effort (Capa et al., 2011; Pessiglione et al., 2007). Considering the frequency with which eye movements can be executed (i.e. 3 saccades per second) the energetic cost of moving the eyes in an environment is rather low. In fact, although the exact cost of eye movements is not known (Shadmehr and Ahmed, 2020), a comparison based on the speed with which other types of movements such as locomotion in an environment or exerting force to a handgrip can be executed (i.e. where each unit of displacement can take several seconds to minutes) suggests that eye movements have lower energetic costs compared to other movement types. This conclusion was supported by studies that simultaneously modelled the energetic costs of eye and hand movements (Diamond et al., 2017). Therefore, an important question is whether an oculomotor task could be challenging enough so that the cost-benefit assessments of effort given rewards are even necessary. Such a task would impose a high enough energy demand for action execution, so that the level of effort is only enhanced when reward is sufficiently high. Here, we devised such a high-demand task, which required the rapid planning of a sequence of saccadic eye movements back and forth between a fixation point and several peripheral targets. Fast and correct execution of the saccades led to the higher magnitude of reward, as the reward gain was scaled with the number of targets landed by the eyes in a fixed amount of time. The level of the reward at stake was briefly shown at the outset of each trial. Importantly, the reward cues were either perceived supraliminally or subliminally, dependent on the relative duration of the mask images that preceded or followed the reward cue (forward and backward pattern masks, respectively). Such a design enabled us to investigate whether the effort entailed by saccadic eye movements is adjusted based on the amount of reward prospect, and whether subliminal and supraliminal reward incentives have similar boosting effects on the saccade performance (Bijleveld et al., 2010; Pessiglione et al., 2007).

We hypothesized that irrespective of their level of conscious awareness, higher reward incentives enhance saccadic effort; measured as the number of targets landed by the saccades as well as frequency and velocity of all saccades in a speeded oculomotor task; as has been shown for manual actions (Pessiglione et al., 2007). Importantly, we tested this hypothesis while stringently controlling for the level of conscious awareness of the reward cues, hence ensuring that subliminal incentives are truly perceived below the level of conscious awareness. Contrary to our hypothesis, we found that subliminal rewards have dissociable effects on different parameters of the saccades. The total number of executed saccades was enhanced by higher rewards irrespective of whether reward cues were perceived subliminally or supraliminally. However, the incentive-dependent adjustment of the peak velocity and accuracy of the saccades seemed to require conscious awareness of the reward magnitude.

## Results

### Visibility test

We manipulated the visibility of reward cues by varying their display duration relative to 2 mask images, one preceding and one following the reward cue (**Figure 1**). Three different display durations were used; the shortest and longest durations were based on a previous study (Pessiglione et al., 2007), whereas the middle duration was set at the detection threshold of each subject (D_indiv_, see Methods and the Supporting Information). To assess the subjective visibility of the reward cues at each display duration, we adapted the 4AFC task used by Pessiglione et al. (2007). In this task, participants choose among 4 options (see 1 cent, see 50 cents, guess 1 cent, guess 50 cents) to report their perception of the coin images. Similar to this study, we assessed both correct responses as well as the number of seen responses, at the group level. This analysis showed that at 100 ms coin images were perceived consciously, as intended. However, the individual display duration, set at the subjective detection threshold for each participant, proved to be unequivocally above the subliminal threshold when assessed with a 4AFC task (for details see the Methods and Supporting Figure 3 and 4). At the lowest display duration (17 ms), the results at the group level were non-conclusive (trend of p= 0.07 for difference from chance level). Therefore, we next applied a more stringent criterion at the individual level. This analysis showed that for 15 participants coins displayed at 17 ms were perceived subliminally (binomial test p>0.05), whereas for 23 participants this display duration was above the level of subliminal perception (binomial test p<0.05). For our subsequent analyses throughout we will refer to the first group as the truly subliminal group and the second group as the supraliminal group (see also the Supporting Figure 4). For each group, the shortest display duration (17 ms) will be referred to as the “low visibility”, the D_indiv_ as the “medium visibility” and 100 ms as the “high visibility” condition.

**Figure 1.**
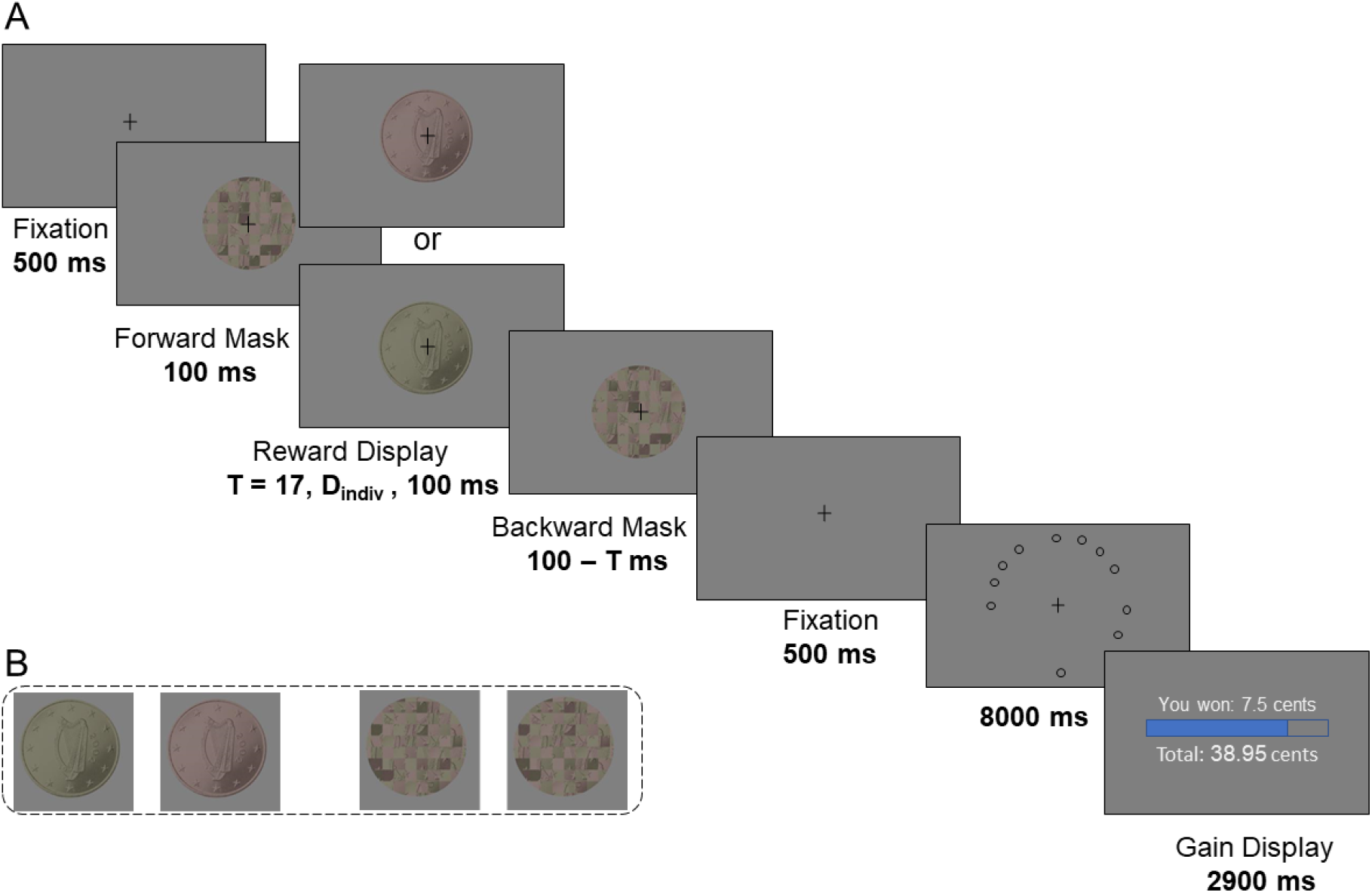
Experimental design. **A. Eye movement task.** After an initial fixation period (500 ms), a sequence consisting of mask-reward-mask images was shown. The reward display was the image of either a 1 cent or a 50 cents coin, shown at one of the three display durations: T= 17 ms, 100 ms or D_indiv_ (an individual display duration between 17 ms and 100 ms, set to the visibility threshold of each subject). After an additional fixation period (500 ms), 18 peripheral targets appeared on the screen at locations randomly chosen on each trial (eccentricity 11.5°). Participants had to look at as many circles as possible in a fixed time (8 seconds). The reward gain was scaled with the number of targets visited by the eyes and was shown at the end of the trial. **B. High and low reward cues and mask images**. From left to right: The 50 cents coin image (high reward cue), the 1 cent coin image (low reward cue), and the complementary checkerboard images that were used as forward and backward masks.

### Visible reward incentives increase the number of hits in a goal-directed oculomotor task

To confirm that our eye movement task (**Figure 1A**) created a demanding situation where the exerted effort is adjusted based on the expected rewards, we first examined participants’ number of hits when coin images were fully visible (100 ms). **Figure 2** illustrates the eye position traces of two example subjects. For each subject, the data of two trials, one with a low and the other with a high reward cue (images of 1 cent and 50 cents coins, respectively) are shown. It can be readily seen that when the amount of expected reward was high, participants made more eye movements and hit more targets compared to when the reward at stake was low. This was the case for participants who had overall lower number of hits (**Figure 2A**), as well as those with higher hit rates (**Figure 2B**). Accordingly, across all subjects (N= 38) the number of hits was significantly higher for high compared to low reward cues (paired t-test, one-tailed, t_37_=2.267, p = 0.01, Cohen’s d = 0.379) (**Figure 2C**). We obtained the same results when a more liberal threshold was applied to count a saccade to the target as a hit (one-tailed paired t-tests, t_37_=2.73, p = 0.004, Cohen’s d = 0.44, also see the Methods). These result show that our paradigm could successfully create a reliable effect of reward incentives on hit rates.

**Figure 2.**
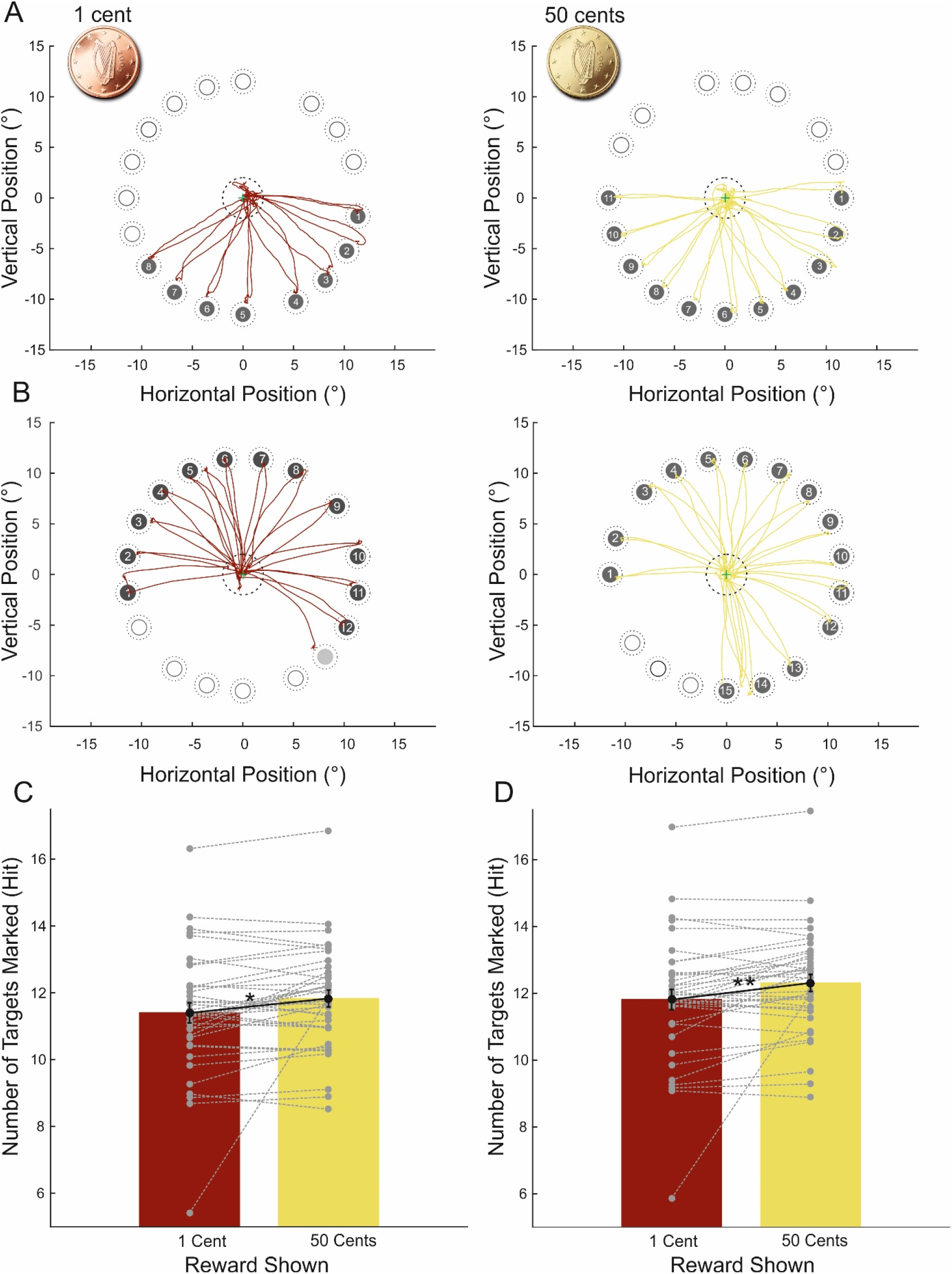
Performance in the eye movement task when rewards were fully visible (displayed at 100 ms). A. Eye position traces of example trials. The data of one example participant in a trial with low (left side, cupper traces) and a trial with high (right side, gold traces) reward is shown. The numbers inside each target indicate the order with which the participant looked at the corresponding target location and the eye movement was counted as a hit. **B** is same as **A**, but was obtained from a different participant who had overall higher number of hits compared to A. The target circle filled with brighter grey color on the left side (low reward condition) was not detected as a hit online, but was regarded as a hit when a more liberal threshold was used offline. **C**. The total number of hits across all participants calculated based on the online threshold (eye position within a 1.2° area from centre of the targets were counted as hits) **D**. same as C but number of hits was calculated based on a more liberal threshold applied offline (eye position within a 2.4° area from centre of the targets were counted as hits). Bars are the average number of hits across participants and small, grey circles correspond to the data of individual subjects. * p<0.05, ** p<0.01. Error bars are s.e.m.

### Comparison of the number of hits across different visibility levels

We next asked whether the incentive-driven effect of reward on hit rates (Figure 2) also extends to other visibility levels where the amount of reward at stake was not fully visible. A mixed-effect ANOVA with participant group (2 levels: subliminal or supraliminal) as between-subjects factor and reward (2 levels: high or low) and visibility (3 levels: low, medium and high) as within-subjects factor was performed on the number of hits. This analysis revealed a main effect of reward as the number of hits was significantly higher for trials with high (50 cents) compared to low (1 cent) incentives (F(1,36) = 4.99, p = 0.032, 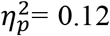). All other main and interaction effects did not reach statistical significance (p>0.05). We obtained similar results when the number of hits was calculated based on a more liberal criterion to count eye movements made towards a target as a hit (**Figure 3**). This analysis revealed a significant main effect of reward (F(1,36) = 8.63, p = 0.006,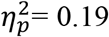) and a significant, but weak interaction between reward and visibility (F(2,72) = 3.76, p = 0.046,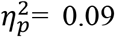). Planned, pairwise comparisons showed a significant effect of reward in all visibility conditions in the supraliminal group (all Ps<0.05, one-tailed paired t-tests) and only at the highest visibility in the subliminal group (t_14_= 2.11, p=0.026, one-tailed, Cohen’s d= 0.54).

**Figure 3.**
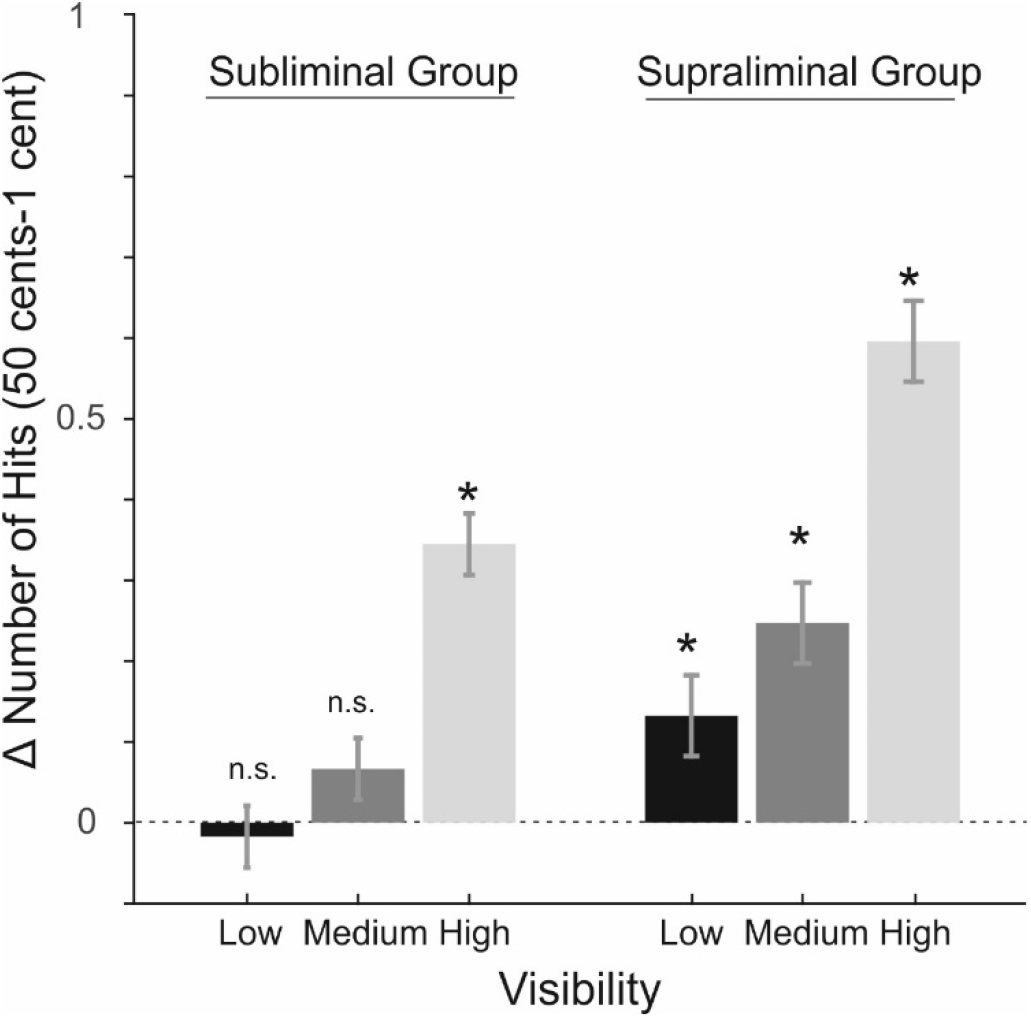
Effect size for the number of hits. Effect size is the difference in the number of hits for high compared to low reward cues (picture of 50 cents and 1 cent respectively). * p<0.05. Error bars are the s.e.m.

### Analysis of the saccade parameters

In order to evaluate the effect of reward incentives on participants’ performance, we have thus far analysed the number of hits. However, the number of hits is a discrete measure that only covers a small subset of saccades that landed adjacent to a target. Therefore, hit rates might not be an accurate indicator of the amount of effort that participants spent, especially when they had aimed to reach a target but failed to do so. We next measured several parameters of the saccadic eye-movements comprising the total number, peak velocity, amplitude and vigor of all saccades, which provide a more sensitive estimation of the amount of spent effort. Additionally, we also measured the landing error of target-directed saccades, characterizing the accuracy of the saccadic plans.

### Reward incentives increase the total number of saccades at all visibility levels

A mixed-effect ANOVA with the participant group as between-subjects factor and reward level and visibility level as within-subjects factors was performed on the total number of saccades (i.e. frequency of saccades within 8 second timeout duration of each trial). This ANOVA revealed a main effect of group (F(1,36) = 6.68, p = 0.014,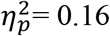) corresponding to higher total number of saccades in the subliminal (mean ± SD = 33.11±3.18 compared to the supraliminal group (mean ± SD = 30.49±2.96, see also Supporting Figure 5B). Importantly, a significant main effect of reward (F(1,36) = 11.85, p = 0.0014,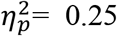) was found. Participants made significantly more saccades in prospect of higher (mean ± SD = 31.84±3.27) compared to lower (mean ± SD = 31.21±3.36) reward magnitudes (**Figure 4A**). All other main and interaction effects were not significant. Planned, pairwise comparisons showed that the effect of reward was significant at all 3 levels of visibility for both groups of subjects (one-tailed paired t-tests, all ps<0.05, all Cohen’s ds> 0.45 in supraliminal group and ds> 0.39 in subliminal group).

**Figure 4.**
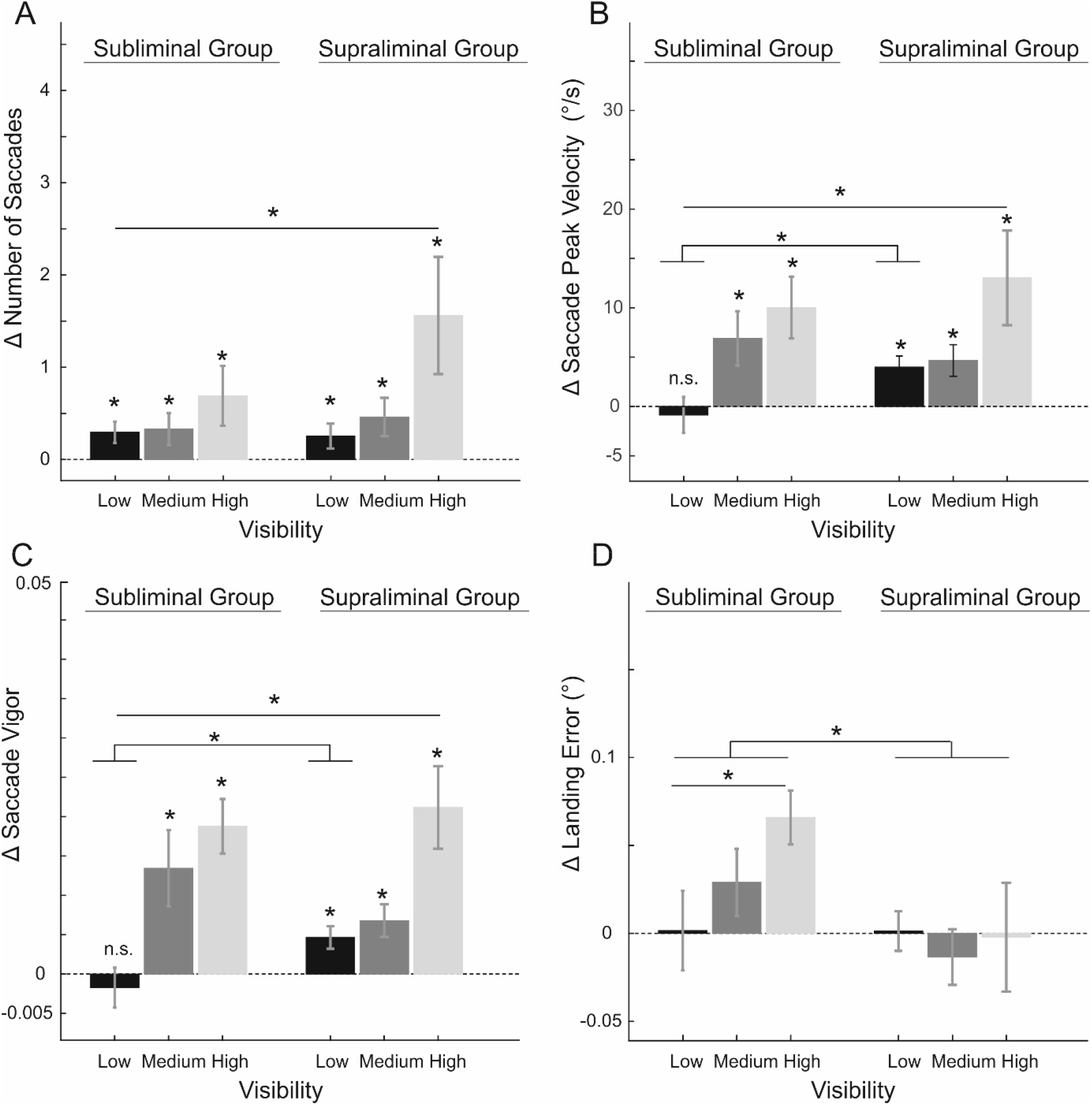
Influence of reward incentives on saccadic parameters. **A**. Effect size for total number of saccades. Effect size corresponds to the difference between 50 cents (high reward) and 1 cent (low reward) conditions in all panels **B**. same as A for the saccade peak velocity **C**. same as A for the saccade vigor **D**. same as A for the saccade landing error. * corresponds to p<0.05. Error bars are the s.e.m.

The main effect of reward and a lack of interaction with other factors indicate that both groups of participants exerted more effort and made more saccades when reward incentives were higher irrespective of their visibility level. This result is in line with previous reports showing that reward incentives can boost effort even when they are perceived subliminally.

### Reward incentives increase the peak velocity and vigor of the saccades only when perceived supraliminally

An important indicator of the performance in the eye movement task is the velocity of each saccade. Optimal adjustment of saccades velocity, through increasing the peak velocity when expected reward is high and decreasing the velocity when rewards are low, ensures that the cost of making a fast saccade is optimally adjusted based on the reward. To investigate whether the peak velocity of saccades are increased by the reward, we performed a mixed-effect ANOVA on the saccadic peak velocity (**Figure 4B**). This ANOVA revealed a significant main effect of reward (F(1,36) = 16.41, p = 0.0002,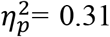) and an interaction between reward and visibility level (F(2,72) = 7.74, p = 0.0035,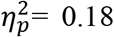) but other main and interaction effects did not reach statistical difference (all ps>0.1).

Since we observed an interaction between reward and visibility level, we performed follow-up mixed-effect ANOVAs at each visibility level. This analysis showed a main effect of reward for the two higher visibility levels (display durations = 100 ms and D_indiv_) that were above the level of conscious perception in both groups (F(1,36) = 12.68, p = 0.0001,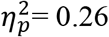 and F(1,36) = 15.15, p = 0.0004,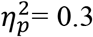, for 100ms and D_indiv_ respectively). However, at the lowest visibility level a significant interaction between group and reward (F(1,36) = 5.79, P = 0.02,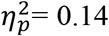) was found. Planned, t-tests at the lowest visibility level revealed that while in the supraliminal group the peak velocity of saccades significantly increased for high compared to low reward incentives (t_22_ = 3.55, p = 0.0008, one-tailed paired t-test, Cohen’s d=0.74), this effect was absent in the subliminal group (t_14_ = 0.48, p = 0.68, one-tailed paired t-test, Cohen’s d=0.12).

Saccadic peak velocity increases in a stereotyped manner when saccade amplitude increases, a relationship known as the *main sequence*. We therefore performed two follow-up analyses to investigate whether the pattern of results observed with the peak velocity of saccades can be explained by possible changes in the saccade amplitude. A mixed-effect ANOVA showed a trend for an increase in saccades’ amplitude when reward incentives were high (F(1,36) = 3.98, p = 0.054,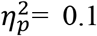). All other main and interaction effects were nonsignificant. Although this analysis did not show a significant effect of reward on saccades’ amplitude, the trend we observed may still indicate that the modulation of saccades’ peak velocity by expected reward partly reflects the influence of reward in increasing saccade amplitudes (hence is a result of main sequence relationship). In a second analysis we tried to remove the effect of saccade amplitude from peak velocities by calculating a within-subject estimate of saccade vigor as done in a previous study (Reppert et al., 2015). This analysis yields a subject-specific ratio between the expected and observed saccade peak velocity for each condition (see Methods). The median of this ratio across all trials of each condition was subjected to a mixed-effect ANOVA (**Figure 4C**). This analysis revealed a significant main effect of group (F(1,36) = 7.34, p = 0.01,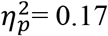), a main effect of reward (F(1,36) = 27.53, p<10^−5^,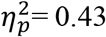) and an interaction between reward and visibility level (F(2,72) = 22.05, <10^− 7^,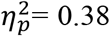). There was a trend for a 3-way interaction between group, reward and visibility level (F(2,72) = 2.86, p = 0.072,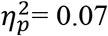) but other main and interaction effects did not reach statistical significance (all p’s>0.1). We next performed follow-up ANOVAs at each visibility level. This analysis showed a significant difference between groups in all visibility levels, with the subliminal group having significantly lower saccadic vigor compared to the supraliminal group (all ps<0.05, all Fs >5, see also Supporting Figure 5D). A significant main effect of reward was found at the two higher visibility levels (F(1,36) = 31.70, p<10^−5^,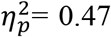 and F(1,36) = 18.8, p= 0.00011,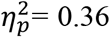 for high and medium visibility, respectively) but at the lowest visibility an interaction between reward and group was observed (F(1,36) = 5.57, p= 0.023, 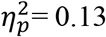). Planned pairwise comparisons at this visibility level revealed that the reward effect was only significant in the supraliminal group but did not reach statistical significance in subliminal group (p= 0.0018, Cohen’s d= 0.67 and p= 0.74, Cohen’s d= 0.17 for supraliminal and subliminal groups respectively, one-tailed paired t-test).

Since we observed a significant effect of group (subliminal or supraliminal) on overall saccade vigor (Supporting Figure 5D), we wondered whether lower vigor in subliminal group is driven by their overall higher uncertainty regarding the magnitude of reward that was at stake on each trial. To test this, we inspected whether across all participants saccadic vigor was correlated with the level of coins’ visibility measured in the 4AFC task (Supporting Figure 6). We found that saccadic vigor was significantly correlated with the level of visibility of reward cues (i.e. the percentage of correct responses) at the shortest display duration (r= 0.46, p= 0.003, Cohen’s d = 1.02). For other 2 visibility levels this correlation was not significant (all ps>0.1).

Thus, the analysis of saccades’ peak velocity and vigor indicate that when reward incentives were presented at a truly subliminal level, i.e. the lowest visibility level in the subliminal group, they did not enhance the saccadic velocity and vigor, while supraliminal rewards showed a strong modulatory effect on both measures. These results indicate that the goal-directed control of saccadic velocity requires conscious awareness. In addition to this effect, we found an overall weaker saccadic vigor in subliminal compared to the supraliminal group. Across all participants (N=38), the level of visibility of reward cues at the shortest display duration was correlated with the within-subject measure of saccadic vigor.

### Reward incentives do not change the speed-accuracy trade-off in supraliminal group but do so in subliminal group

The performance in the eye movement task depends on both speed (i.e. the number of all saccades and the velocity of each saccade) and accuracy of saccades in landing on the target. We found a main effect of reward on the number of saccades and velocity of each saccade across all participants. At the lowest visibility level however, the effect of reward was only significant in the supraliminal group. If the increase of saccade frequency and velocity occurs without concomitant increase in saccadic landing error, the reward-driven invigoration of eye movements does not occur at the cost of a decrease in their accuracy (i.e. speed-accuracy trade-off). To test this, we next examined whether the saccadic error, i.e. distance of saccades’ endpoints from the targets, was changed with reward incentives at different visibility levels and in the two group of subjects (**Figure 4D**).A mixed-effect ANOVA with participant group as between-subjects factor and reward and visibility level as within-subjects factors was performed on the saccade’ endpoint errors. This ANOVA revealed a significant interaction effect between group and reward (F(1,36) = 4.62, p = 0.038,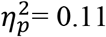). There was a trend for a main effect of group (F(1,36) = 3.83, p = 0.058,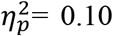) but this effect did not reach statistical significance (see also Supporting Figure 5E). All other main and interaction effects were non-significance. The interaction effect between group and reward was further investigated by performing post-hoc pairwise comparisons for the effect of high versus low reward, separately in each group of participants. In the subliminal group the endpoint error was significantly larger when reward incentives were higher (t_14_= 3.28, p= 0.003, Cohen’s d=0.84, two-tailed paired t-test). However, in the supraliminal group higher rewards did not lead to a higher saccadic error (t_22_= 0.39, p = 0.65, Cohen’s d= 0.08, paired t-test, two-tailed). These results suggest that in the subliminal group the control of saccades’ speed-accuracy is not optimal and an increase in saccades’ frequency and velocity happens at the cost of a decrease in their accuracy. This pattern was not observed in supraliminal group in line with previous reports showing that reward incentives increase both speed and accuracy of saccades (Manohar et al., 2015). One explanation for this effect is that subconscious perception of reward magnitude at the shortest display interval triggers the subliminal group to change their overall criterion for an optimal control of speed-accuracy when faced with higher rewards. If this is the case, across all subjects (from both groups) the reward-driven change in saccade landing errors should be negatively correlated with the level of coins’ visibility at the shortest display duration (Supporting Figure 7). We found that this was in fact the case, as a negative correlation (r = −0.33, p= 0.042, Cohen’s d=0.7) was found between the visibility reports at the 17 ms and the reward effect (i.e. endpoint error in high reward minus endpoint error in low reward trials) across all subjects (N=38).

### The specificity of the reward effect for target-directed saccades at different levels of visibility

Do the reward incentives specifically boost saccades traversing the distance between the target and the fixation point; as warranted by the goal of the task? To answer this question, we carried out an exploratory analysis, where we inspected the reward effect on saccades’ peak velocity (Figure 4B) as a function of its amplitude. To this end, we first examined the probability distribution of saccade amplitudes pooled across all subjects (**Figure 5A**). As expected, this distribution contained two peaks: one (the highest) corresponding to the saccades with an amplitude that was close to the distance between the target and the fixation point (i.e.11.5°). These saccades included large eye movements that traversed the distance between the fixation point and the target (i.e. back or forth between the two). Another smaller peak was found at amplitudes <2°, most likely corresponding to corrective saccades bringing the eyes to the centre of the target or the fixation point. Remarkably, the effect of rewards on saccades’ peak velocity nicely matched the spatial profile of the probability distribution of saccade amplitudes, being maximum at the amplitudes matching the target eccentricity (**Figure 5B-D**). Importantly, while the reward effect was broadly distributed across almost all saccade amplitudes for fully visible reward cues (**Figure 5B**), it showed a narrower distribution centred at larger saccades that fulfilled the goal of the task (i.e. to move the eyes between the target and the fixation point) at lower visibility levels (**Figure 5C-D**). Crucially, for the lowest visibility level (17 ms), reward affected the velocity of the target-directed saccades only for supraliminal group and only for the saccades that matched the eccentricity of the target (**Figure 5D**). The spatial specificity of the reward effect suggests that at lower visibility levels, the uncertainty ensuing from subconscious perception of rewards is compensated by narrowing down rewards’ boosting effect to only those eye movements that could maximally aid the pursuit of the task goal. However, this optimization process only happens if reward incentives are consciously perceived.

**Figure 5.**
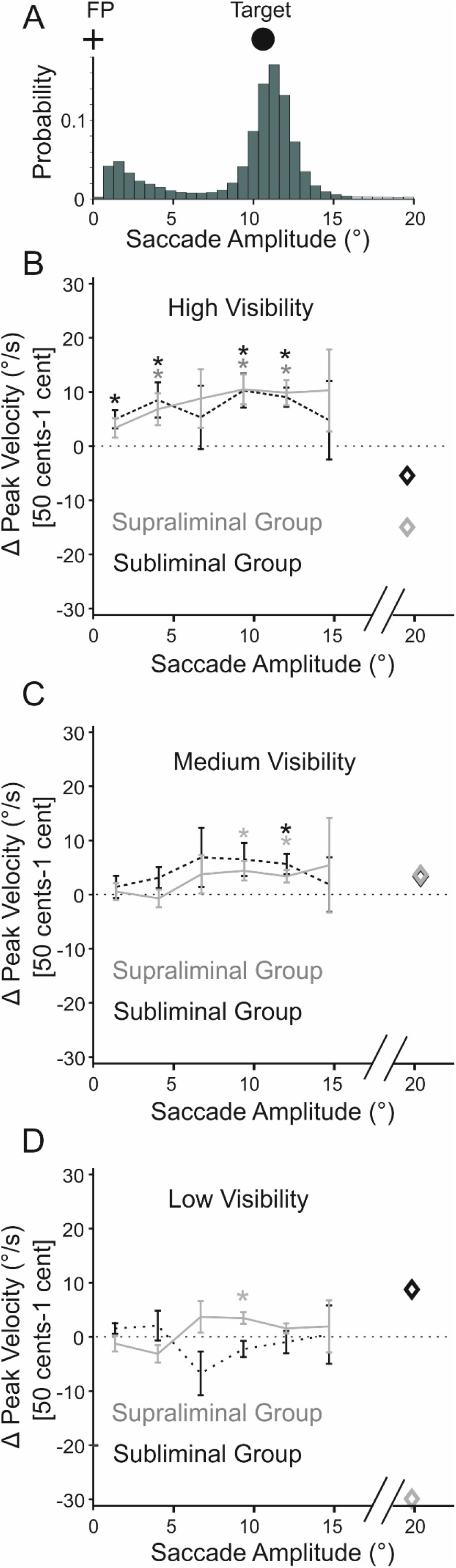
The effect of reward incentives on saccade velocity specifically occurs for target-directed saccades, but only when perceived consciously. **A. Probability distribution of the saccade amplitudes**. The peaks of the distribution correspond to the target eccentricity (11.5°, schematically illustrated by the distance between the black circle; i.e. target; from the fixation point: FP) and the corrective saccades (peak at <2°). Saccades with eccentricities >16° occurred <0.1% percent of time and are highlighted with the light green color at right tail of the histogram **B-D. The spatial profile of the reward effect for different visibility levels of reward cues. B**. For fully visible cues the reward effect occurs at almost all saccade amplitudes and for both groups of participants (supraliminal: grey and subliminal: black). **C**. At medium visibility reward effect in both groups is only significant for saccades that have an amplitude close to the target-fixation distance. **D**. At the lowest visibility level, reward effect is solely significant for saccades with an amplitude equal to the target-fixation distance, and only for the supraliminal group. In panel B-D diamond markers are the average peak velocities for all saccades >16°. Error bars are the s.e.m. Stars correspond to significant effects at p<0.05, based on two-tailed paired t-tests.

### Results obtained with the analysis of saccade parameters are independent of the saccade detection method

We have so far used a fixed velocity threshold (35°/s) to detect saccades. We next tested whether we obtain similar results when saccades are detected based on an adaptive algorithm (Nyström and Holmqvist, 2010). The rationale for this analysis was to rule out that the pattern of results we observe are because of differences in the trial-by-trial level of noise in eye movements across different conditions and especially between the two groups of participants.

Specifically, the lack of a significant effect on the velocity and vigor of saccades despite an increase in the total number of saccades at the lowest visibility level in the subliminal group could be due to higher level of noise in the eye position data when rewards are high, for instance due to an overall enhancement of saccades velocity due to the arousal enhancing effects of reward. The adaptive algorithm introduced by (Nyström and Holmqvist, 2010) can be applied to the data of individual trials of each subject, therefore being adjusted for different levels of noise across participants and conditions. Applying this analysis to the eye position data, we found overall similar results as when a fixed velocity threshold was used to detect saccades (see Methods and The Supporting Information for details).

## Discussion

Previous studies have shown that reward incentives could enhance the exerted effort in prospect of a higher reward, even when presented subliminally (Bijleveld et al., 2009; Pessiglione et al., 2007; Zedelius et al., 2014). The aim of the current study was to test whether these effects extend to the eye movements. We observed an enhancement of participants’ oculomotor effort; including number of hits and frequency, velocity and accuracy of all saccades; when higher reward was at stake and reward cues were fully visible, as expected. Subliminal rewards on the other hand only increased the total number of saccades but they did not enhance saccades’ peak velocity/vigor. Additionally, participants who had reached the truly subliminal level of perception at the shortest visibility level showed an overall decrement in accuracy as a function of reward, as across all visibility levels saccade endpoint errors were larger when higher rewards were at stake. This suboptimal speed-accuracy trade-off did not occur in the supraliminal group. Examination of the strength of reward effect across saccade amplitudes revealed that the boosting effect of reward specifically occurs for saccades to or from the target (i.e. with an amplitude that matched the target-fixation point distance), especially at lower visibility levels. This boosting effect however did not happen for truly subliminal rewards. These results therefore demonstrate a dissociation between the effects of subliminal rewards on saccades’ initiation and control of velocity and accuracy.

The results obtained with the fully visible reward cues (i.e. 100 ms) are in line with previous studies showing a modulatory effect of reward incentives on saccade metrics (Chen et al., 2014; Choi et al., 2014; Hikosaka et al., 2000; Takikawa et al., 2002). Specifically, we found an increase in the number of saccades and their velocity/vigor when higher rewards were at stake and reward cues were perceived consciously. This finding indicates that humans can voluntarily adjust their oculomotor performance by changing the number of eye movements they make and the velocity and vigor of them depending on the magnitude of expected rewards. As such, these results confirm previous findings that saccade metrics could be, at least to some extent, voluntarily controlled (Chen et al., 2014; Muhammed et al., 2020). Importantly, the invigorating effect of supraliminal rewards also occurred when the stereotypical relationship between the saccade amplitude and velocity was factored out, as shown by the within-subjects measure of saccade vigor. However, the energizing effect of rewards on saccade metric could also be driven by involuntary mechanisms, as rewards have been shown to exert automatic/reflexive modulation of saccade metrics (Manohar et al., 2017). The conjoint involvement of voluntary and involuntary mechanisms is supported by theoretical frameworks suggesting two distinct influences of reward motivation on behavior (Dickinson and Balleine, 1994; Niv et al., 2006). According to these, goal-directed effects determine the current goal of behavior (e.g. obtaining food or water when hungry or thirsty), and affect behavior in an ‘outcome-specific’ manner. In contrast, ‘energizing’ effects of reward are more general and determine the vigor of all actions, not only those that are related to current goals but also other pending or pre-potent actions (Dickinson and Balleine, 1994; Niv et al., 2006). In other words, energizing effects of reward m a y occur in an ‘outcome-unspecific’ and involuntary manner (but see also Manohar et al., 2017). As such the motivational effect of reward that we found in the current study may be a combination of both of voluntary and involuntary mechanisms.

While we found similar effects of reward incentives on saccade initiation (i.e. total number of saccades produced in a fixed amount of time) and velocity when reward cues were fully visible, a dissociation was found for subliminal rewards. In participants who had reached a subliminal level of perception at the lowest visibility level, higher rewards increased the total number of saccadic eye movements but did not increase the average peak velocity of the saccades. In addition, we also found that the saccadic endpoint error was not optimally adjusted for high reward incentives in subliminal group across all visibility levels. The observation that the number of produced saccades is enhanced by reward incentives, regardless of their level of consciousness, is in line with previous studies showing an effect of subliminal cues (Bargh et al., 2001; Kay et al., 2004) and specifically subconscious reward incentives (Aarts et al., 2008; Bijleveld et al., 2012, 2009; Capa et al., 2013, 2011; Holland et al., 2005; Pessiglione et al., 2007; Zedelius et al., 2014) on goal-directed, effortful behavior. However, the lack of an effect of subliminal rewards on saccade velocity/vigor contradicts these previous findings and instead supports other studies which showed a divergence, either behaviorally or at the level of brain responses, between the motivational effects of conscious and subconscious reward cues (Bijleveld et al., 2014, 2010; Correa et al., 2018; Zhan et al., 2017). Whereas some of the latter studies suggested that this divergence occurs very under specific cognitive conditions, as for instance in self-versus others-judgements (Zhan et al., 2017), others showed that such divergence is more generic and occurs whenever a strategic adjustment of behavior is needed. This was exemplified in situations where only conscious compared to subconscious reward incentives resulted in an optimal speed-accuracy trade-off (Bijleveld et al., 2010), adjustment of performance based on the attainability of the expected rewards (Zedelius et al., 2012), and learning to modify choices depending on the outcomes (Correa et al., 2018). To understand our results in light of these previous findings, we note that by initiating more saccades participants could increase their total number of hits in the task we employed. However, the increase in the number of hits and hence the obtained rewards also depended on an optimal adjustment of the velocity and accuracy of each saccade. We found that neither the velocity nor the accuracy of the saccades were optimally adjusted when reward cues were subliminal, which supports the proposal that optimal speed-accuracy adjustment in speeded tasks requires conscious awareness (Bijleveld et al., 2010).

What can explain the observation that saccade initiation could be steered by subconscious reward signals, whereas the control of saccade velocity and accuracy requires conscious awareness of incentives? We posit that motivational influences on saccades’ initiation and their control of velocity and accuracy may rely on separable neural mechanisms. Although this proposal cannot be directly tested in our current eye tracking study, several lines of evidence from previous studies support our conjecture. Burst activity of neurons in Superior Colliculus (SC) underlies saccade initiation (Sparks, 1978; Wurtz and Goldberg, 1971). Additionally, microstimulation (Katnani and Gandhi, 2012; Stanford et al., 1996), and inactivation (Lee et al., 1988) studies have demonstrated that SC neurons are involved in adjustment of saccades’ peak velocity. Therefore, the Superior Colliculus could potentially enhance both the initiation and the velocity of the saccades. Collicular neurons however, receive inputs from a large number of cortical and subcortical structures involved in processing of reward information (for reviews see Hikosaka, 2007; Hikosaka et al., 2014). Among the subcortical inputs, projections from the Substantia Nigra pars reticulata (SNr) and Caudate Nucleus (CD) to SC play an important role in modulation of saccade initiation based on reward (for reviews see Hikosaka, 2007; Hikosaka et al., 2014, 2000), as both structures (CD and SNr) receive input from dopaminergic neurons (Hikosaka et al., 2014). The Cortical inputs to SC, on the other hand, comprise excitatory inputs from the frontal eye field (FEF), Supplementary eye field (SEF) and lateral intraparietal area (LIP). These areas, for instance the FEF, are an important source of top-down information to the SC (Wurtz et al., 2001), play a role in the control of the voluntary saccades (Dias et al., 1995; Gaymard et al., 1998), and are sensitive to the changes in reward magnitude (Chen and Wise, 1996; Glaser et al., 2016; Platt and Glimcher, 1999). Importantly, these cortical areas, particularly the FEFs, provide inputs to the Caudate Nucleus (CD), thereby indirectly (through CD-> SNr ->SC) affecting saccade initiation and control.

Together these findings from previous neurophysiological research indicate that purposeful eye movements are controlled by a distributed network encompassing subcortical and cortical brain areas. This network tightly interacts with reward-related brain areas in basal ganglia (BG) and particularly CD and SNr. The subcortical part of this network encompassing CD-> SNr ->SC pathway can initiate and control saccades adjusted for reward magnitude and hence may underlie the effect of subliminal rewards on saccade initiation that we observed. However, the additional inputs from cortical areas plays a paramount role in optimal voluntary control of saccades dependent on the reward magnitude. The lack of optimal adjustment of saccades’ peak velocity and accuracy based on reward magnitude for subliminal incentives may indicate that the involvement of the cortical supervisory control of saccades requires conscious awareness. As indicated by our results regarding the spatial specificity of the reward effects (Figure 5B-D), at lower visibility levels an adaptive mechanism specifically limits the reward modulation to the target-direct saccades. This finding has similarity to the findings of neurophysiological studies where the activity of cortical and subcortical areas were simultaneously recorded while an animal was engaged in a reward learning tasks (Ding and Hikosaka, 2006). It was shown that while both cortical and subcortical areas, exemplified by the FEF and Caudate nucleus, contribute to the reward-related biases in generation of the saccades, FEF played a more important role in optimizing the accuracy of a saccade whereas CD was more prominent in coding of reward value. Hence cortical pathways involved in saccade generation and control may be part of a network that exerts supervisory control over oculomotor plans depending on the specific task demands and external or internal feedbacks (Coe et al., 2019; Munoz, 2002), and they do so only if the task demands are consciously appraised. This tentative proposal can be experimentally tested by future neuroimaging studies.

The idea of separable mechanisms for the motivational control of saccade initiation and velocity and accuracy adjustment is further supported by findings from clinical studies (for a review see Leigh and Kennard, 2004). For instance, in Parkinson’s disease (PD), characterized by loss of dopaminergic neurons in the Substantia Nigra pars compacta (SNpc), the execution of goal-directed, volitional saccades is impaired, while reflexive saccades have normal metrics (Lal and Truong, 2019). This impairment is ascribed to disruption of the frontal cortex-BG-SC circuit mentioned before. An elegant recent study (Manohar et al., 2015), tested whether saccade abnormalities in PD are a result of a general problem in economic evaluation of the reward information that could ensue from such disruption (Shadmehr et al., 2019). To test this idea, a speeded saccade task was used where reward depended on the saccade latency of voluntary saccades. In control group, reward increased saccadic velocity and endpoint accuracy, and reduced RTs. Interestingly, in the PD group these effects occurred to a much lesser degree, which is in line with previous reports of suboptimal saccade control in these patient (Rand et al., 2000). Computational modeling revealed that this impairment could be modelled by assuming a greater cost of controlling noise. Importantly, the PD patients had no problem in initiation of the saccades as the total number of saccades was equal in both groups. These results suggest that the disruption of the frontal cortex-BG-SC circuit as exemplified in PD may specifically impairs goal-directed saccadic control while leaving saccade initiation, elicited by neurons downstream to the Basal Ganglia (BG) intact.

While previous studies have investigated how different levels of conscious awareness of saccade goals affect the eye movements, our study is the first to investigate the effect of subconscious reward incentives on goal-directed saccades. In one previous study (van Zoest and Donk, 2010), saccadic selection as a function of the degree of awareness of a saccade goal was examined. Observers performed a visual search task and made an eye movement to a specific item in the display. The display was masked contingent on the first eye movement of the subjects. This study showed that participants’ saccadic performance was completely unrelated to their awareness of the saccade goal, as performance was identical regardless of whether participants could report specific features of the saccade target. A more recent study showed that participants have a gaze bias towards a stimulus that they are completely unaware of its position and identity (Rothkirch et al., 2012). Finally, in a free viewing task, it was shown that visual cues presented pre-saccadically and below the awareness threshold could influence the direction and latency of the first free saccade (Huang et al., 2014). Based on these findings it has been suggested that saccadic selection is primarily driven by subconscious processes. This idea is further supported by a more general theoretical framework proposing a dissociation between action and perception (Milner and Goodale, 2008), accounting for cases where participants can be unaware of the perceptual features of a saccade target, but could nevertheless execute flawless actions towards them. Our results partially support these proposals but also show important differences, as we demonstrate that some aspects of saccade planning could be steered by subconscious information, whereas others such as optimal control of saccade velocity and accuracy require conscious awareness.

Another related line of research has investigated the effect of motivational signals on conscious and subconscious eye movements (Blaukopf and DiGirolamo, 2006). This study employed an antisaccade task where trials were either rewarded for a correct response or punished for errors. In addition to the antisaccade task, participants indicated on each trial whether they were aware of the accuracy of their performance. Although high valence trials of both positive and negative rewards led to the slowing down of antisaccades, unconscious errors were instead speeded when punishment was high. Speeded responses have in general lower accuracies. Thus when punishment is high, the best strategy would be to slow down the saccades so that higher accuracies are reached. Instead, this study showed that when participants were unaware of their errors, they failed to maintain an optimal speed-accuracy tradeoff. Our results are in line with this suggestion and further support the idea that optimal adjustment of actions based on motivational signals requires conscious awareness.

We propose that our results demonstrate a general dissociation between motivational influences on the initiation and velocity control of saccades, with the latter requiring conscious awareness. However, future research is needed to elucidate whether the pattern of results we obtained is a general characteristic of motivational effects on eye movement planning or rather they depend on the specific features of the task we employed. One aspect of the task we used is particularly important in this respect: participants needed to plan a sequence of eye movements towards multiple targets. In a previous study, it was shown that when humans are required to execute saccades to a large number of target locations, saccade preparation for all target locations is carried out in parallel (McSorley et al., 2019). It is hence possible that in our experiments, subjects who had perceived rewards subliminally differed in the way they planned a sequence of saccades compared to subjects who were consciously aware of the rewards. In fact, the strategy used for planning saccades to a sequence of targets may discourage careful planning of individual saccades in favor of increasing the rate of saccadic production, as has been shown previously (Wu et al., 2010). This explorative strategy emphasizes on producing more secondary corrective saccades to compensate for saccadic landing errors of primary target-directed saccades, particularly when the target locations are peripheral and adjacent to each other (Wu et al., 2010). This pattern has a striking similarity to what we observed in subliminal subjects, as they had higher number of saccades as a group compared to the supraliminal subjects, while they had lower accuracy of primary target-directed saccades. A tentative proposal is that the conscious awareness of rewards changes the balance of favored strategies when a sequence of eye movements is planned, a proposal that can be tested in future studies.

Although, a wealth of previous research has demonstrated that goal-directed behavior could be controlled by subconscious processes, a great deal of controversy surrounds the reported effects (for a review see Newell and Shanks, 2014). One important concern regarding subliminal effects is the uncertainty regarding whether participants had truly reached a subconscious threshold. Typically, researchers try to overcome this difficulty by showing that subliminal cues are not perceived above chance on a group level. However, this method does not consider the cases when a group consists of a mix between individuals who had reached the subliminal threshold and those who had not. One important strength of our study is the strict measure for subliminal perception at the individual level, where each person was evaluated with respect to the level of awareness of reward cues, as opposed to majority of previous studies that used a criterion at the group-level to determine subconscious perception of reward. This allowed us to avoid falsely supporting that subliminal perception occurred in all individuals as a group. Additionally, by ascertaining which individuals belonged to either the truly subliminal or the supraliminal group, we were able to elucidate differences in oculomotor planning that occurred in each group separately, which would have otherwise been impossible (e.g. the overall change in number of saccades in subliminal compared to supraliminal group). One may argue that the strength of the masks we employed was suboptimal, hence preventing all individuals to reach the subliminal threshold. While this is a possibility that we cannot rule out, we note that our sample size was large enough to allow having sufficient number of individuals belonging to each group and hence we could draw robust conclusions regarding the differences and similarities of reward effects across the two groups.

Taken together, our results suggest that some aspects of reward-driven oculomotor planning could be modulated by subconscious processes, whereas others need conscious awareness. Our study underscores the importance of understanding oculomotor planning as a means to gain insight into underlying mechanisms of cognition. Saccades play a pivotal role in acquiring sensory information across space and are closely linked to attentional selection. Previous studies have shown that saccade metrics could be used to probe different aspects of decision making in both perceptual (Seideman et al., 2018) as well as value-based decision making (Shadmehr et al., 2019). Our study furthers these previous findings and additionally shows that saccades provide detailed and precise information regarding how one of the most complex aspects of human cognition, namely that of conscious awareness, is orchestrated in the brain.

## Materials and Methods

### Participants

Forty subjects (20 male and 20 female, age 19 to 45 years; mean ± std age, 25.65 ± 5.04 years) participated in the experiment for financial compensation. All but six were right handed and had normal or corrected-to-normal vision, and were naïve to the hypothesis of the project. Two participants were excluded from the final analysis: for one participant part of the data was lost due to the technical problems during the experiment and for the other participant more than 25% of all trials in the eye movement task had to be removed (see also Methods: Data Analysis). Thus, the final sample comprised 38 subjects.

Before the experiment started and after all procedures were explained, participants gave their oral and written consent. The study was approved by the local ethics committee of the “Universitätsmedizin Göttingen” (UMG), under the proposal number 15/7/15.

### Stimulus presentation and eye tracking Apparatus

Throughout the experiment, visual stimuli were displayed on a calibrated ASUS monitor subtending 1280 x 800 pixels, and a refresh rate of 120 Hz placed at a distance of 60 cm to the participants. For tracking the eye position an EyeLink 1000 Plus system with a desktop mount was used (SR Research). The monocular eye position data (right eye) was acquired at a rate of 1000 Hz for all but three participants. Due to a technical error the data from these participants (N=3) was recorded at a rate of 250 Hz and was interpolated to a sampling rate of 1000 Hz during the offline analysis. All experiments were scripted in MATLAB, using Psychophysics toolbox (Brainard, 1997). The EyeLink camera was controlled by the corresponding EyeLink toolbox in MATLAB (Cornelissen et al., 2002). Before each block, the eye tracking system was calibrated to provide precise measurements (using a 13-point standard EyeLink calibration procedure).

### Eye Movement Task

We employed a speeded eye movement task with a 2 (Reward: 50 cents vs 1 cent) by 3 (Display Duration: 17ms, D_indiv_= an individual display duration, 100ms) within-subjects factorial design. In this task, participants were exposed to the picture of a coin with either 50 cents (high reward) or 1 cent (low reward) value at the beginning of each trial (**Figure 1**). They could earn a fraction of the shown reward by making a sequence of eye movements from the fixation point towards 18 peripheral target circles. In some trials the coin image was displayed for 100 ms, allowing subjects to clearly discriminate the coin value (supraliminal), whereas in other trials shorter durations of either 17 ms or an individual display duration (D_indiv_, set individually at the visibility threshold of each subject), were employed. The longest and shortest presentation durations (100 ms and 17 ms) were adapted from a previous study that used a similar masking procedure (Pessiglione et al., 2007). The individual duration (D_indiv_) was determined through a staircase procedure (see also the Supporting Information). The eye movement task started with a practise block of 12 trials followed by six blocks of 42 trials each. This comprised 42 trials for each condition, counterbalanced for two values of the coin image (1 and 50 cents) presented at three display durations.

The sequence of events in a trial was as follows: a trial began with a fixation period (500 ms, fixation area 1°). Thereafter a forward mask (100 ms) followed by the picture of a coin (1 cent or a 50 cents Euro coins, displayed at 3 different durations: 17 ms, D_indiv_ or 100 ms), and a backward mask (100 ms minus coin display duration) were displayed while the participant was required to maintain fixation on the central fixation cross overlaid on the picture of the masks and the coin. Subsequently, a second fixation period (500 ms) was followed by the main eye movement task (**Figure 1**). During the eye movement task, 18 circular targets (radius 0.75°, eccentricity 11.5°) surrounded a fixation cross at the centre of the screen. In each trial, the position of the targets (N=18) was selected by randomly drawing 18 samples out of 42 possible locations. This was done to discourage participants from planning their sequence of eye movements prior to the presentation of the targets (since target locations in each trial were unpredictable, also see Figure 1). Participants started the task by looking at the fixation cross and subsequently made an eye movement towards a target at any location. A target was considered to be “*marked*” (the term used for subjects; i.e. a hit) as soon as the eye position fell within a circular area with a radius of 1.2° from the centre of the target. In our offline analysis (Figure 2C), we counted a target-directed eye movement as a hit, if the eye position fell within an area twice as large as the threshold used online (radius = 2.4°). Once a target was landed, it got filled with black color, thus indicating to the participant that they had successfully hit the target. A landed target remained black for the rest of the trial. In order to land on the next target, the participants had to first focus back on the fixation cross. The color of the fixation cross turned from black to red whenever the subject refocussed the fixation cross (within an area of 2.5° from the centre of the fixation cross). This way, the participant received immediate feedback and was informed that they could continue to look at the next circle. As the participant moved their eyes away from the fixation cross, it turned black again. The total duration of the eye movement task was fixed to 8 s on each trial.

The goal of the participants in this task was to land on as many targets as possible in a fixed time interval (8 s). Depending on how many target a subject could hit on each trial, they received a certain fraction of the coin’s value displayed at the outset, calculated as:

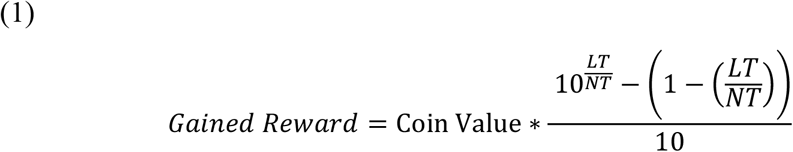

Where *LT* is the number of landed targets, *NT* corresponds to the total number of targets, and Coin Value is either 1 or 50 cents. The exponential equation was used to potentiate the effect of incentives, as participants could readily see that their number of hits had a strong influence on the gained rewards. The gained reward was visually displayed on the monitor at the last phase of each trial and stayed in view for 2900 ms. Here, participants could see how much money they gained during the last trial and also the amount they had earned so far in the current block of the experiment. Additionally, a reward bar was shown to illustrate the progress of the reward earning during a block of the experiment. This bar was scaled to the total maximum amount that a participant could earn during a block of the experiment.

### Visibility Tests

#### I. Staircase procedure to determine the individual display duration D_indiv_

We used a one-up and one-down staircase method (Kaernbach, 1991; Wetherill and Levitt, 1965) to determine the threshold at which an individual could no longer identify the coins (for details see the Supporting Material).

#### II. Four-alternative forced choice (4AFC) task

We employed a “4-alternative forced choice” (4AFC) task adapted from a previous study (Pessiglione et al., 2007) to assess the perceptual visibility of the reward cues at each display duration (i.e. 17 ms, D_indiv_ and 100 ms). To assess participants’ level of conscious perception of reward at each display duration, we averaged the performance (correct responses of either “seen” or “guess”) across the 2 repetitions of the 4AFC task (once before and once after the eye movement task, for details see the Supporting Information).

### Data analysis

The data obtained from all parts of the experiment was analysed using custom-written scripts in MATLAB (version R2015a).

#### I. Analysis of the visibility tests

In order to assess whether participants had reached subliminal perception at the shorter display durations, the data from both 4-AFC before and after the eye movement task was pooled and analysed. This comprised a total of 80 trials at each display duration. We first analysed the data following the approach used by Pessiglione et al. (2007), where subliminal perception needs to meet two criteria: Firstly, correct answers (either “seen” or “guess”) should not differ from chance level. Secondly, “seen” responses – independent of correct or incorrect – should not be different from 0. To test for these criteria, one-sample t-tests were used to test whether “correct” or “seen” answers at a group level were different from chance (50%) or 0, respectively. Chance-level performance at a group level in such a forced-choice discrimination task is a typical criterion to decide whether participants have reached subliminal perception (Overgaard and Sandberg, 2012; Sandberg et al., 2010). This analysis showed that at 100 ms and the individual detection threshold (D_indiv_) correct responses were significantly higher than the chance level (mean ± SD = 99.5 ± 0.92%, t_37_= 52.12, p<10^−10^, Cohen’s d =8.34 and mean ± SD = 84.6 ± 13.8%, t_37_= 1.47, p = 0.017, Cohen’s d =0.4 respectively, one-tailed, one-sample t-test). At the shortest display duration (17 ms) correct responses were not significantly different from chance, albeit a trend was found (mean = 61± 7%, t_37_= 1.47, p= 0.075, Cohen’s d=0.24, one-tailed, one-sample t-test). Analysis of the “seen” responses showed that they were not significantly different from 0 at the shortest display duration (mean ± SD = 10 ± 13%, p =0 .219, one-tailed, one-sample t-test), but they significantly differed from 0 at the other two display durations (mean ± SD = 98 ± 01.9%, and 44±12%, for 100 ms and D_indiv_ respectively, all ps<10^−3^, one-tailed, one-sample t-test). These results indicate that the coin images were perceived consciously at 100 ms, as intended. However, the individual display duration set at a level where participants subjectively reported not seeing the coins proved to be above the level of conscious awareness when tested by a 4AFC task. The results at the lowest display duration were not conclusive, as according to one of the criteria (i.e. the “seen” responses) coin images were perceived subliminally while the correct responses showed a trend suggesting that for some participants this display duration has been longer than the threshold for subliminal perception.

We therefore used a more stringent criterion at the individual level to test whether the shortest display duration was truly perceived subliminally. To this end, performance of each individual at 17 ms was tested against chance performance using a binomial test (one-sided implemented in MATLAB). According to this tests subjects who had up to 47 out of 80 correct responses were assigned to the subliminal group (see Supporting Figure 4). Subjects who had more than 47 out of 80 correct responses were sorted into the supraliminal group. Using this criterion 15 participants were classified as truly subliminal, whereas 23 were categorized as supraliminal (total sample size N=38).

#### II. Analysis of the eye movement task

To assess different saccade parameters, eye position data of each trial from the beginning to the end of the eye movement task was analyzed (total duration 8000 ms). Trials with >25% missing data in this period were discarded from the analysis. To detect a saccades, eye position samples in a trial were smoothed using a Savitzky-Golay low-pass filter with an order of 2 and length of 20 ms (Nyström and Holmqvist, 2010). Saccade onsets were defined as the moment when a sample exceeded a two-dimensional velocity threshold of 35°/s. Saccade offsets were calculated as the first sample where the eye position velocity and acceleration dropped below 35°/s. Additionally, saccades with an inter-saccadic interval shorter than 40 ms and a duration shorter than 10 ms were discarded from the analysis.

We also tested whether we obtain similar results when saccades were detected based on an adaptive algorithm (for details of the results see Supporting Information) as introduced by Nyström and Holmqvist (2010). This algorithm can be applied to the data of individual trials where the threshold to detect saccades is set based on the noise-level of each trial (hence the threshold is adaptive and not fixed, accounting for variability of noise level across trials and subjects). Furthermore, since the local noise level of samples are taken into account when the start and end of the saccade are estimated, the adaptive method is especially suitable for situations when participants scan the visual scene by making multiple eye movements, as is the case in the paradigm we employed here. We used the adaptive algorithm as described in (Nyström and Holmqvist, 2010) while setting the peak threshold equal to the mean plus 4 standard deviation of the trial velocity, saccade onset threshold equal to the mean plus 2.5 standard deviation of the trial velocity and the minimum saccade duration at 10ms. Saccades with preceding fixation durations shorter than 40 ms were discarded from the analysis (Nyström and Holmqvist, 2010).

For both methods of saccade detection, the reported values of peak velocity and vigor (see below) were measured based on the maximum two-dimensional velocity of the samples between the onset and the offset of the saccade. Saccade amplitude was calculated as the Euclidian distance between the eye position samples at the onset and offset of the saccade.

We computed a within-subject measure of saccade vigor, as shown in Figure 4 and Supporting Figure 5 and 6, using a method employed in a previous study (Reppert et al., 2015). To this end, we measured the amplitude (represented by *x*) and the peak velocity (represented by *v*) of all saccades across all trials of each participant. Subsequently a hyperbolic function in the following form was fit to the saccade amplitude data:

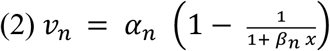

Where 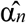 and 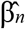 characterize the hyperbolic relationship between the peak velocity and amplitude of the saccades. Based on these parameters, for each participant an expected saccade velocity given each saccade amplitude was calculated 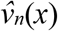 Note that fit parameters and hence the expected velocity is computed from all saccades of a particular individual and hence solely reflects the stereotypical biomechanical relationship between velocity and amplitude captured known by the main sequence (Bahill et al., 1975). Finally, the ratio between the measured and the expected velocities was calculated as the median of 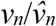 across all trials of each condition (i.e. reward level and display duration). We used median rather the mean of 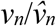 to reduce the impact of potential outlier trials. This ratio represents a within-subject measure of saccade vigor with ratios >1 reflecting a greater than average vigor in a certain condition in an individual.

To quantify saccades’ accuracy, we first determined whether a saccade was directed towards the peripheral targets or the fixation point. Subsequently, we measured the distance of the endpoint of the saccade from its respective destination as the saccadic landing error. Target-directed saccades were defined as saccades that either landed < 5.75° (i.e. half of the targets’ eccentricity = 11.5°) from the peripheral targets or moved the eyes away from the fixation point and towards the peripheral targets (i.e. their endpoint was farther from the fixation point than their start point) while landing >5.75° away from the targets. The remaining saccades were categorized as fixation-directed. For saccades that their endpoint was more than 11.5+5.75° from the fixation point, the minimum distance was set to the minimum distance from any target irrespective of whether the saccade was going towards or away from the fixation point.

For statistical inferences, we analyzed the number of hits and the saccadic parameters (number of saccades, peak velocity and vigor and endpoint error) by carrying out a series of mixed-effect, repeated measures analyses of variance (ANOVAs). The ANOVAs included one between-subjects factor (subliminal or supraliminal group based on the results of the binomial test on 4AFC task) and two within-subjects factors: *reward* (2 levels: 1 or 50 cent) and *visibility level* (3 levels: low, medium and high corresponding to 17 ms, D_indiv_ and 100 ms display durations) as independent factors. To correct for violations of the sphericity assumption, a Greenhouse–Geisser correction was applied to the degrees of freedoms and p-values of all the ANOVAs. Significant effects from the omnibus ANOVAs were further investigated by performing paired t-tests (one-tailed). Using one-tailed tests was warranted since based on previous studies (Pessiglione et al., 2007), we hypothesized a directional difference between high and low reward conditions with respect to these parameters (i.e. higher rewards enhance saccadic parameters compared to low rewards). However, we did not have a hypothesis regarding the impact of reward on saccades’ endpoint error and hence applied two-tailed paired t-tests for pairwise analyses. Effect sizes in ANOVAs are reported as partial eta-squared 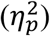 and in pairwise comparisons as Cohen’s d (Lakens, 2013) throughout.

## Acknowledgements

We thank Adem Saglam for his help with programming of the experiments, Danilo Postin for his help with the analysis of the eye position data and Christiane Valuch and Thorsten Albrecht for their valuable comments on the analysis methods. This work was initially supported by a seed fund grant from Leibniz Science Campus *Primate Cognition, Göttingen, Germany* to MW and AP and continued by an ERC Starting Grant (no: 716846) to AP.

## Authors’ contributions

MW and AP conceptualized the project and acquired funding. OU, VKH and AP designed the task. VKH and OU conducted the experiments. VKH, OU and AP analyzed the data. VKH, OU, JEA, MW and AP interpreted the results and wrote the first draft of the manuscript. All authors revised the manuscript.

## Supporting Information

### Supporting Material

#### Participants

Our participants were invited via an online recruiting system (http://www.probanden.eni-g.de/orsee/public/). The experiment took about 1 hour and 45 minutes in total. Participants were told that they could earn between 10 to 25 Euros according to their performance. After the experiment, participants received the amount of money they earned during the experiment in cash.

#### Reward cues and masking procedure

During the visibility tests and the eye movement tasks participants had to discriminate whether the displayed coin image had a value of 1 cent (low reward) or 50 cents (high reward). For this purpose, Irish 50 cents coin images were obtained from www.muenzen.eu and processed with an image editor. This processing included changing the luminance of the image by half and subsequently transforming one of the coin images to the other. To this end, the image of the 50 cents coin was used as a template and its hue in HSV colour system was changed from red to yellow to obtain a 1 cent coin. This was done to ensure that coin images are identical in all but one respect (colour), hence preventing any visual bias that could result from a difference in the structure of the coin images (**Figure 1B**).

As the forward and backward masks, we used patterned checkerboard images consisting of both 1 cent and 50 cents image fragments. To create a checkerboard, square-shaped regions (referred to as tiles, size = 72 pixels, n = 38 tiles per coin) from each of the two coins were randomly selected and repositioned alternately over the area of the mask image. Moreover, to avoid similarity in the structure of the masks and the coins, tiles were selected in a way that neighbouring regions from the same coin were avoided (at least 3 neighbouring tiles were skipped). Additionally, a different checkerboard, built by re-randomization of the tile orders, was used in each trial for the forward and backward masks to prevent a flicker (resulting from rapid succession of identical image elements). The reasoning for this was that participants could use a perceived flicker between the forward mask, coin image, and backward mask as a hint towards which coin has been displayed (to report seeing the coin with a colour that did not flicker). Hence, to avoid this confound a forward mask and a distinct complementary backward mask were created for each trial (see **Figure 1B**). This way the complementary mask does not create a flicker when participants look at a certain tile or region, since there is only one change in colour during the mask-coin-mask pattern presentation.

#### Sequence of tasks during an experimental session

In each session, participants first performed a short discrimination task to get familiar with the coin images. For this, images corresponding to either 1 cent or 50 cent coins were displayed for variable durations. After a coin was displayed, the participant was asked to decide which coin they had seen. After 16 trials, the accuracy was calculated based on the participant’s correct responses. Each coin was shown 8 times with the 8 different display duration without any masking, ranging from 17 – 100 ms. In case a participant could not distinguish between the pictures of the coins with an accuracy of at least 80%, no further testing was performed. After the discrimination task, each participant had to perform the staircase, followed by the 4AFC tasks (see below). Thereafter, the main eye movement task and finally the second 4AFC task were conducted. Between blocks of different tasks participants could rest and the experiment was resumed after recalibration of the eye tracker.

### Visibility Tests

#### I. Staircase procedure to determine the individual display duration D_indiv_

We used a one-up and one-down staircase method (Kaernbach, 1991; Wetherill and Levitt, 1965) to determine the threshold at which an individual could no longer identify the coins (Supporting Fig.1). Here, the order of events in each trial were similar to the eye movement task but after the second fixation period participants were asked whether they could see a coin or not. Participants were instructed to only report seeing the coin when they could clearly identify which of the two coins (1 or 50 Cents) were displayed. They responded with the right arrow of the keyboard for “I saw a coin” and with the left arrow for “I did not see a coin”. Based on their response, the display duration of the next trial was adjusted to one of 7 possible values (i.e. D = 25, 34, 42, 50, 59, 67, 100 ms; selected to be multiples of the monitor’s frame rate). The display duration of the first trial was set to 100 ms and was subsequently decreased by one step if the participant reported seeing the coin. Likewise, if the participant reported not seeing the coin, the display duration of the next trial was increased by one step. To obtain the individual visibility threshold for each participant (D_indiv_), we calculated the mean of the last five turning points of the staircase and subtracted one frame duration from it if it was >25 ms (see Supporting Figure 2 for the distribution of D_indiv_).

#### II. Four-alternative forced choice (4AFC) task

We employed a “4-alternative forced choice” (4AFC) adapted from a previous study (Pessiglione et al., 2007) to assess the perceptual visibility of the reward cues at each display duration (i.e. 17 ms, D_indiv_ and 100 ms). The shortest (17 ms) and the longest (100 ms) durations were the same as in Pessiglione et al. (Pessiglione et al., 2007) and were intended to produce either a subliminal (17 ms) or a supraliminal (100 ms) perception of the reward incentives. The middle duration (D_indiv_) was individually adjusted for each participant to be at the border of subliminal perception. To assess participants’ level of conscious perception of reward at each display duration, we averaged the performance (correct responses of either “seen” or “guess”) across the 2 repetitions of the 4AFC task (once before and once after the eye movement task). The 4AFC task was structured in the same way as the staircase and eye movement task (Supporting Figure 3): a trial began with a fixation period (500 ms) followed by the presentation of a forward mask (100 ms), the image of either a 1 cent or a 50 cents coin (17, D or 100 ms), a backward mask (100 ms minus coin display duration), and a second fixation period (500 ms). Participants subsequently chose among 4 options to indicate which coin they saw or guessed to have seen. To do so, they could either press “A” or “S” keys indicating “seen 50 cents” or “seen 1 cent”, or press “D” or “F” keys indicating “guess 50 cents” or “F” for “guess 1 cent” respectively. All conditions were counterbalanced, with 20 repetitions per incentive value and reward timing.

### Results obtained with using an adaptive algorithm to detect the saccades

We also tested whether we obtain similar results when saccades were detected based on an adaptive algorithm as introduced by Nyström and Holmqvist (2010). This algorithm can be applied to the data of individual trials where the threshold to detect saccades is set based on the noise-level of each trial (hence the threshold is adaptive and not fixed, accounting for variability of noise level across trials and subjects). Using this analysis we found overall similar results to the ones reported in the main text. In brief, higher reward incentives were associated with higher number of saccades across all visibility levels and both participant groups (F(1,36) = 13.10, p = 0.0008, 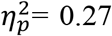 for the main effect of reward) and subliminal group made overall more saccades than the supraliminal group (F(1,36) = 5.98, p = 0.019,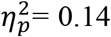 for the main effect of group). Analysis of the saccadic vigor showed a significant main effect of reward (F(1,36) = 28.5, p<10^−5^,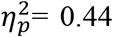 for the main effect of reward) and an interaction between reward and the display duration (F(2,72) = 21.55, p<10^−7^, 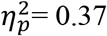). Also a trend was found for a significant 3-way interaction between reward, display duration and group (F(2,72) = 2.47, p = 0.094,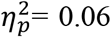). Follow-up analyses showed that saccades’ velocity and vigor were not affected by reward incentives at the lowest visibility level in subliminal group (p = 0.56 and p = 0.75, for both velocity and vigor, paired t-test, one-tailed, Cohen’s d= 0.04 and d=0.18 respectively) but higher rewards led to higher saccadic peak velocity and vigor in all other comparisons (all ps<0.05, all Cohen’s ds>0.55). The comparison of saccades’ endpoint error showed a significant effect of group (F(1,36) = 4.38, p= 0.043,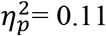) corresponding to larger landing errors in subliminal group, but the interaction between group and reward did not reach statistical significance (p>0.1). Taken together, the results of the adaptive saccade detection method showed similar results to the fixed threshold but also slight differences were found. These differences could be due to the fact that the thresholds used for detection of the saccades was set at the level of individual trials and therefore corrected for effects that were driven by an overall difference across individuals but were unrelated to within-subject modulations by reward.

## Supporting Figures

**Supporting Figure 1.**
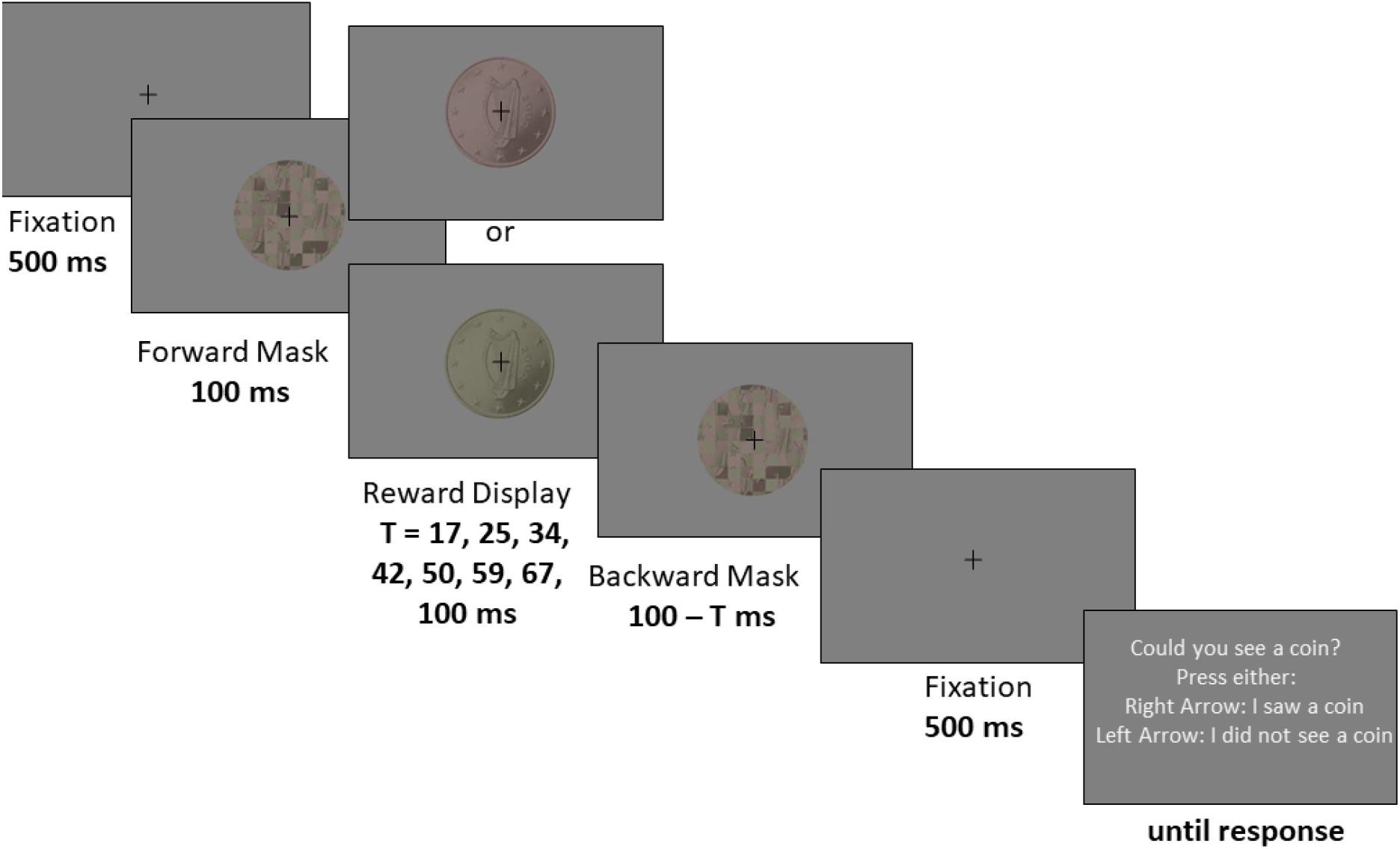
Experimental design of the staircase procedure used to determine the detection threshold of each participant (D_invdiv_). After an initial fixation period (500 ms), a sequence consisting of mask-reward-mask images was shown. The reward display was the image of either a 1 cent or a 50 cents coin, shown at an individual display duration between 17 ms and 100 ms (steps increasing by 8 ms, i.e. the refresh rate of the monitor). The display duration was changed based on participants’ reports, being increased one level when they reported not seeing the coin and decreased one level when they saw the coin. The mean of the last 5 turning points of the staircase was calculated to determine the D_invdiv._

**Supporting Figure 2.**
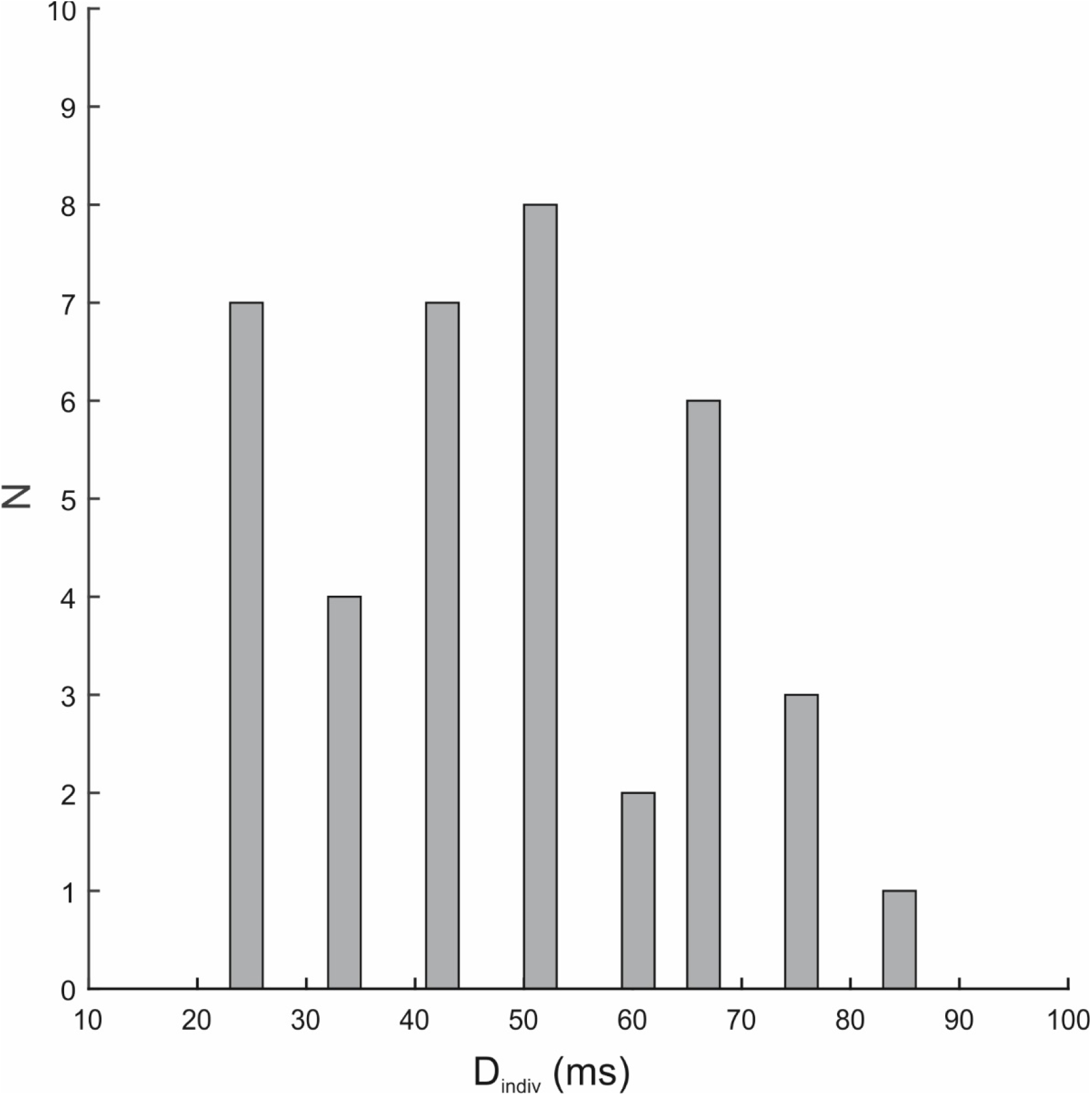
Distribution of detection threshold (D_invdiv_) across all participants.

**Supporting Figure 3.**
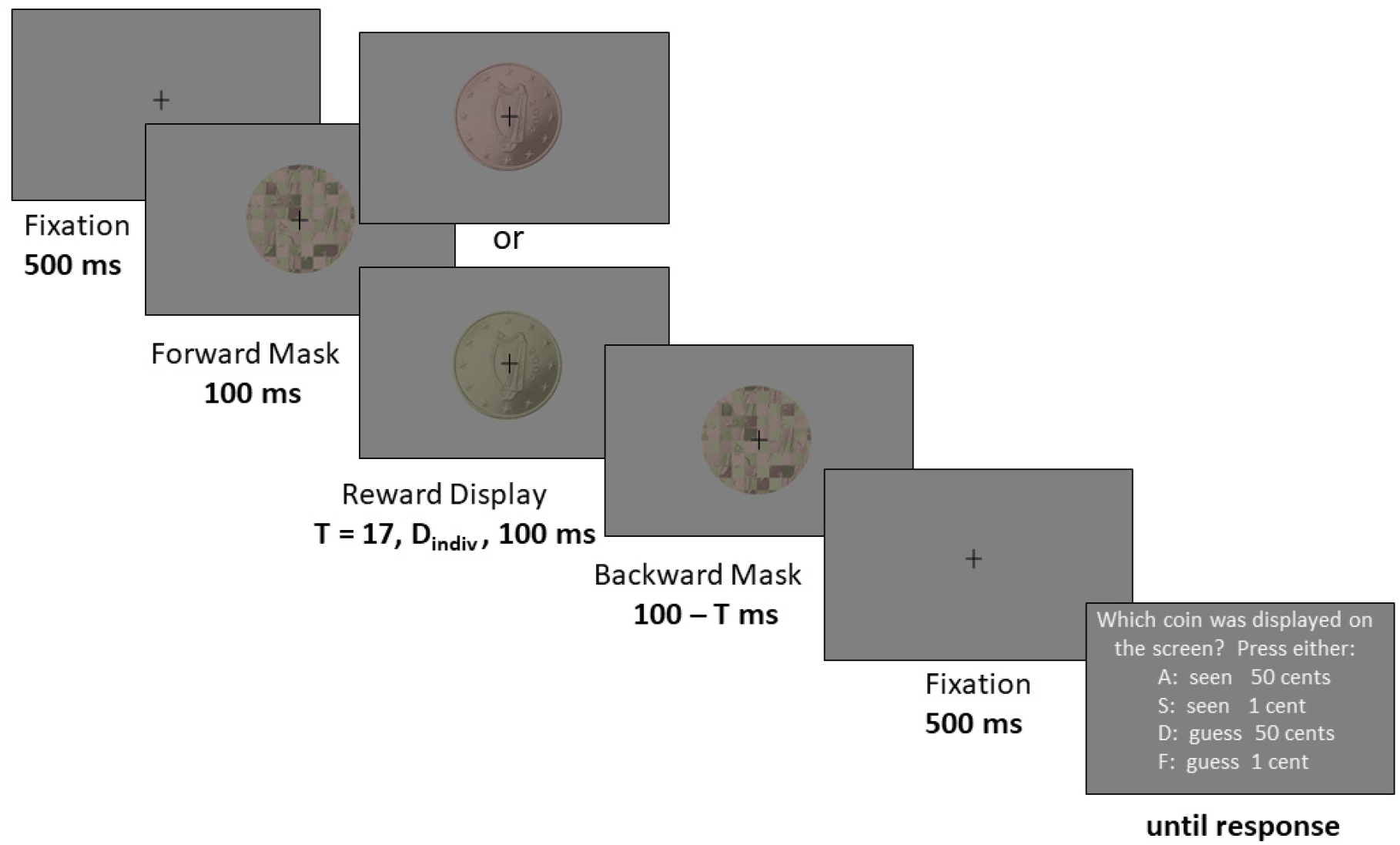
Experimental design of the 4AFC task. After an initial fixation period (500 ms), a sequence consisting of mask-reward-mask images was shown. The reward display was the image of either a 1 cent or a 50 cents coin, shown at one of the three display durations: T= 17 ms, 100 ms or D_indiv_ (an individual display duration between 17 ms and 100 ms, set to the visibility threshold of each subject). The 4AFC task had the same task structure as the eye movement task except that after the fixation-reward-coin-mask-fixation sequence, participants were asked about their perception of the displayed reward magnitude. The 4-alternative-forced-choice (4AFC) task was used to test the level of visibility of the reward cues at each display duration, and was done once before and once after the eye movement task.

**Supporting Figure 4.**
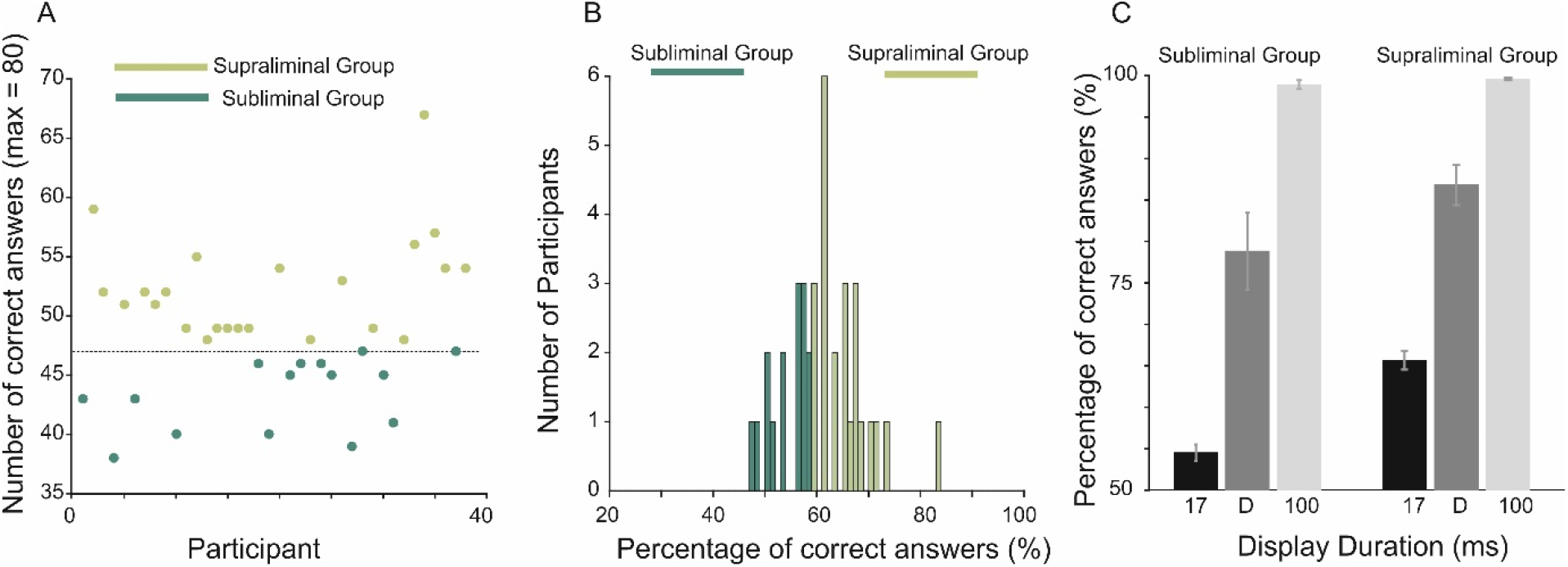
Results of the 4AFC visibility test. **A**. Based on a binomial test if correct responses are than 47 out of 80 trials (i.e. the dashed line), the performance is significantly higher than chance level. Based on this test performed on the data of the shortest display duration (17 ms), participants were divided to two groups: subliminal participants had correct responses (seen+guess) lower or equal to 47, whereas in supraliminal participants correct responses were more than this cut-off. **B**. Percentage of correct responses are shown for the two group of participants that were divided based on criterion shown in A, at the shortest display duration. **C**. Probability of correct identification of the reward value is shown separately for subliminal and supraliminal groups at each visibility level. Error bars are the s.e.m.

**Supporting Figure 5.**
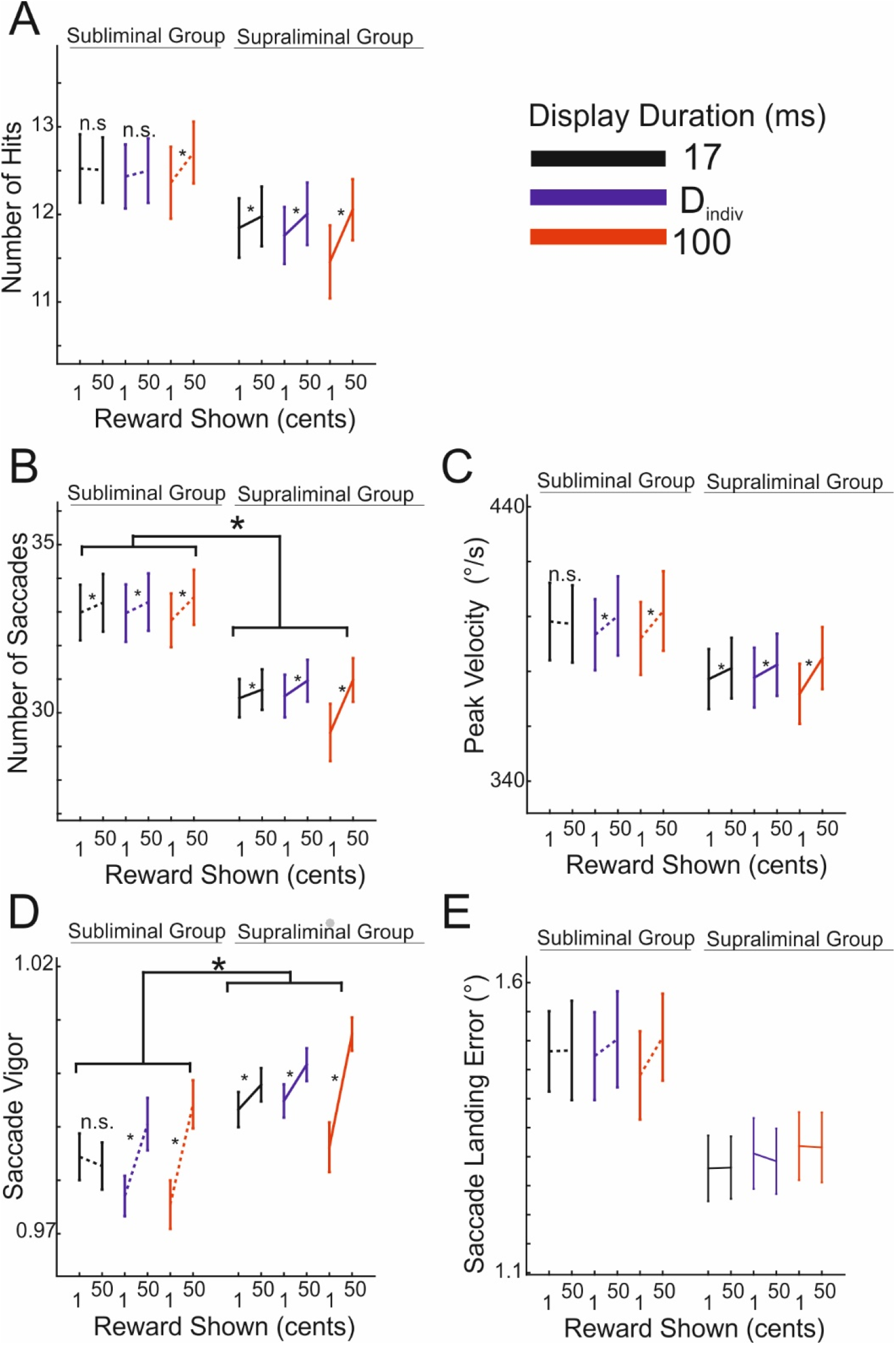
Performance parameters for each reward and visibility level. This figure is complementary to Figure 4 in the main text. Here, instead of the reward effect size (high-low reward), the net mean value of each parameter is shown. This allows to compare differences in overall magnitude of a certain parameter across visibility levels and between participant groups (subliminal versus supraliminal). To ease the comparison with Figure 4 in the main text, only the effect of reward based on pairwise comparisons at each visibility level (i.e. display duration) are highlighted. In addition, significant effects of participants’ group are illustrated. **A**. Hit rates. **B**. Total number of saccades (subliminal group > supraliminal group) **C**. Saccade peak velocity **D**. saccade vigor (subliminal group < supraliminal group) **D**. saccade landing error. * corresponds to p<0.05. Error bars are the s.e.m.

**Supporting Figure 6.**
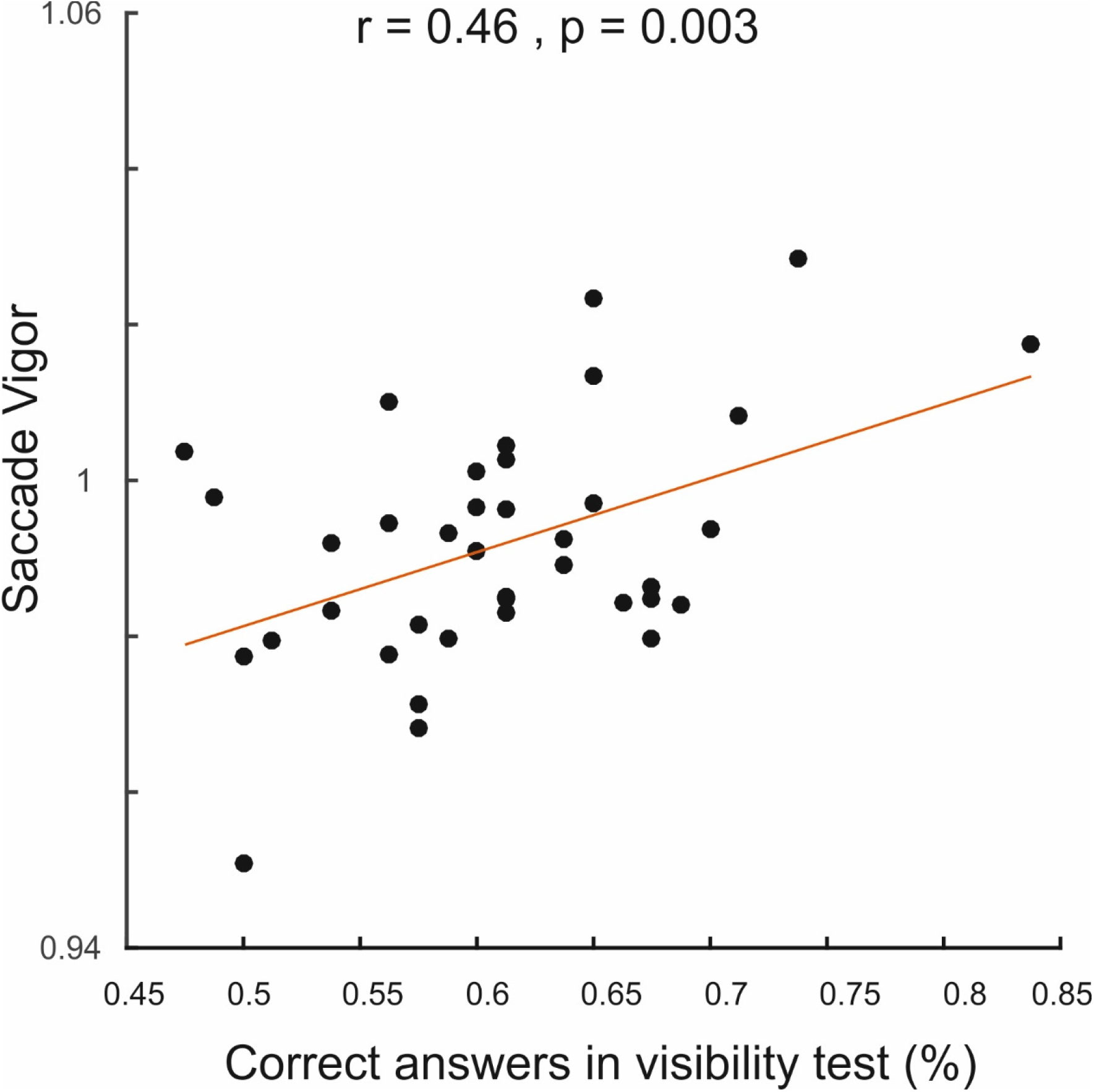
Correlation of the average saccadic vigor with the visibility reports at the shortest display duration. Each point represents the average vigor across all conditions in one individual.

**Supporting Figure 7.**
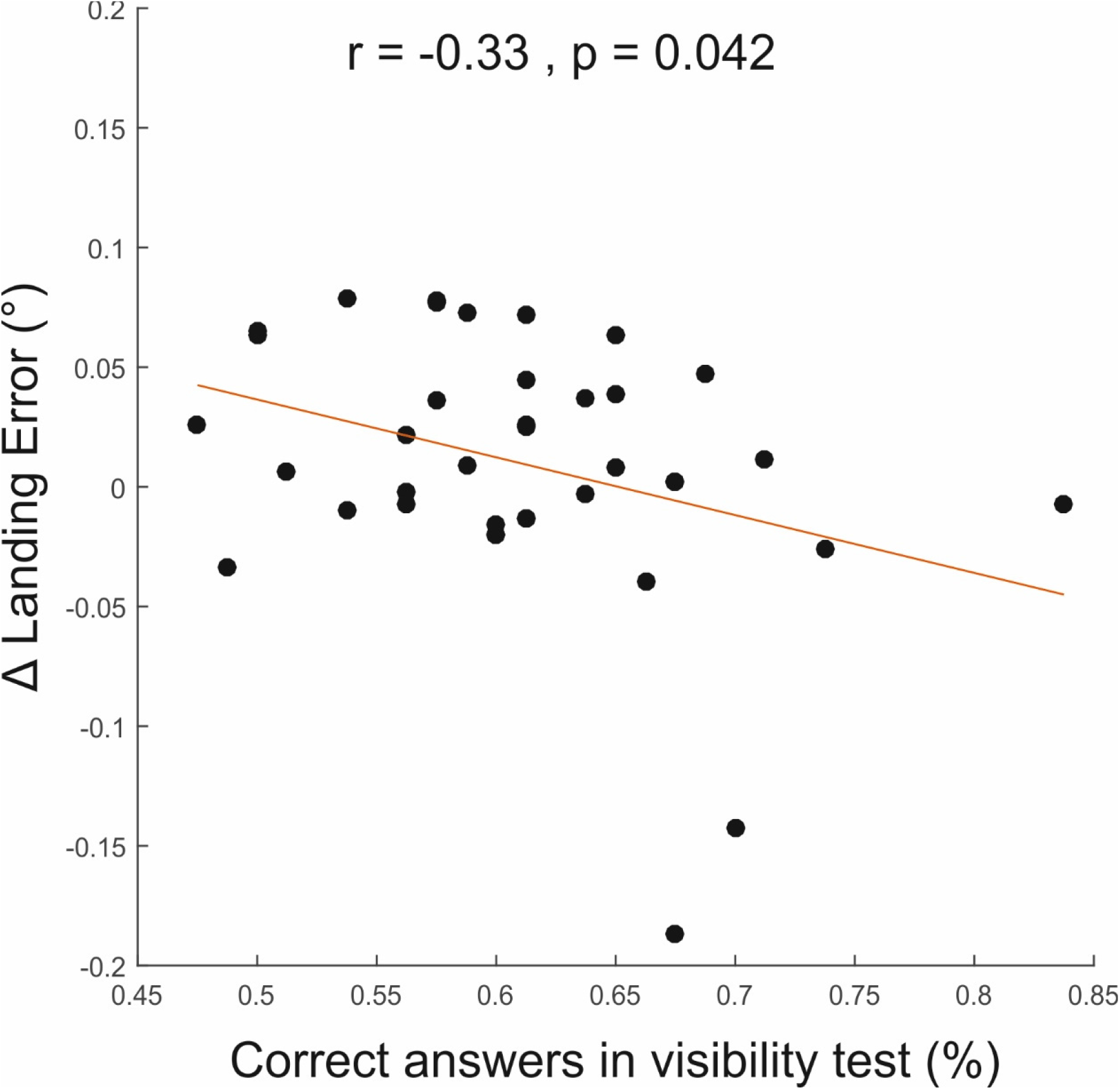
Correlation of the reward effect on landing errors (high-low reward across all visibility levels) with the visibility reports at the shortest display duration. Each point represents the average effect size across all display durations in one individual.

